# Transient “rest” induces functional reinvigoration and epigenetic remodeling in exhausted CAR-T cells

**DOI:** 10.1101/2020.01.26.920496

**Authors:** Evan W. Weber, Rachel C. Lynn, Kevin R. Parker, Hima Anbunathan, John Lattin, Elena Sotillo, Zinaida Good, Meena Malipatlolla, Peng Xu, Panos Vandris, Robbie G. Majzner, Yanyan Qi, Ling-Chun Chen, Andrew J. Gentles, Thomas J. Wandless, Ansuman T. Satpathy, Howard Y. Chang, Crystal L. Mackall

**Author notes:** Correspondence: Crystal L Mackall MD, Lorry Lokey Stem Cell Bldg. 265 Campus Drive, Ste 3141A, MC5456 Stanford, CA 94305.

## Abstract

T cell exhaustion limits immune responses against cancer and is a major cause of resistance to CAR-T cell therapeutics. Using a model wherein tonic CAR signaling induces hallmark features of exhaustion, we employed a drug-regulatable CAR to test the impact of transient cessation of receptor signaling (i.e. “rest”) on the development and maintenance of exhaustion. Induction of rest in exhausting or already-exhausted CAR-T cells resulted in acquisition of a memory-like phenotype, improved anti-tumor functionality, and wholescale transcriptional and epigenetic reprogramming. Similar results were achieved with the Src kinase inhibitor dasatinib, which reversibly suppresses CAR signaling. The degree of functional reinvigoration was proportional to the duration of rest and was associated with expression of transcription factors TCF1 and LEF1. This work demonstrates that transient cessation of CAR-T cell signaling can enhance anti-tumor potency by preventing or reversing exhaustion and challenges the paradigm that exhaustion is an epigenetically fixed state.

## INTRODUCTION

Chimeric antigen receptors (CARs) combine a tumor antigen-recognition domain with intracellular signaling domains, thereby enabling recognition and killing of tumor cells in a major histocompatibility complex (MHC)-independent manner (Lim and June, 2017; Mackall et al., 2014). CAR-T cells mediate high response rates in relapsed/refractory high-grade B cell malignancies, but less than 50% of patients experience long-term disease control and CAR-T cells have not demonstrated reproducible efficacy against solid tumors (Davila et al., 2014; Lee et al., 2015; Majzner and Mackall, 2019; Maude et al., 2014). T cell exhaustion has been implicated as an important factor limiting the efficacy of CAR-T cells against cancer (Eyquem et al., 2017; Fraietta et al., 2018a; Fraietta et al., 2018b) and can be driven by excessive CAR signaling as a result of high antigen burden or tonic signaling induced by antigen-independent clustering of the CAR receptor (Long et al., 2015; Lynn et al., 2019). We hypothesized that transient cessation of CAR signaling would enable exhausted CAR-T cells to regain functionality and form a memory pool, similar to the effects observed following antigen clearance of acute infections (Wherry et al., 2004).

Using a drug-regulatable platform (Banaszynski et al., 2006; Banaszynski et al., 2008) wherein a tonically signaling CAR was modified with a C-terminal destabilizing domain (DD) to enable drug-dependent control of CAR protein levels, we observed that transient inhibition of CAR surface expression, and thereby tonic CAR signaling (i.e. “rest”), prevented cells from developing exhaustion, instead redirecting them to a memory-like fate with enhanced anti-tumor potency. Similarly, transient rest in a population that had already acquired functional, transcriptional and epigenetic hallmarks of exhaustion, but not *α*PD-1 blockade, induced impressive functional reinvigoration that was associated with global transcriptional and epigenetic reprogramming. We observed similar phenotypic and functional reinvigoration following transient exposure to dasatinib, a Src kinase inhibitor that potently and reversibly inhibits TCR and CAR signaling (Lee et al., 2010; Mestermann et al., 2019; Schade et al., 2007; Weber et al., 2019). Collectively, these results challenge the paradigm that exhaustion is an epigenetically fixed state (Pauken et al., 2016; Philip et al., 2017) and reveal that transient cessation of CAR signaling may provide a new strategy for augmenting the function of exhausted human CAR-T cell populations.

## RESULTS

### A GD2-targeting CAR modified with a destabilizing domain exhibits drug-dependent, analog control of expression and function *in vitro* and *in vivo*

We previously demonstrated that some CARs, like those incorporating a GD2-specific 14g2a scFv and CD28 co-stimulatory domain, undergo antigen-independent, tonic CAR signaling due to spontaneous receptor clustering, which promotes hallmark features of exhaustion in human T cells (Long et al., 2015; Lynn et al., 2019). To test whether prevention of tonic signaling during *ex vivo* expansion would preserve CAR-T cell functionality and improve anti-tumor potency, we incorporated an FK506 binding protein 12 (FKBP) destabilizing domain (DD) (Banaszynski et al., 2006) into the endodomain of the GD2-targeting CAR (GD2.28*ζ*.FKBP). The DD induced rapid proteasomal degradation of the CAR fusion protein at baseline (Figures 1A and 1B), whereas in the presence of shield-1, which specifically binds to and stabilizes the FKBP DD, CAR degradation was prevented and the GD2.28*ζ*.FKBP CAR was expressed on the cell surface in a dose-and time-dependent manner (Figures 1B and 1C). Removal of shield-1 from GD2.28*ζ*.FKBP-expressing T cells lead to a rapid decrease in CAR surface protein, with a degradation half-life of approximately 1 hour (Figure 1C). To determine whether the dynamic range of CAR expression induced by the DD was sufficient to modulate biologic reactivity and to directly assess the relationship between CAR expression level and anti-tumor function, we co-cultured GD2.28*ζ*.FKBP CAR-T cells with Nalm6 leukemia engineered to overexpress GD2, GFP, and luciferase (Nalm6-GD2) in the presence of increasing concentrations of shield-1. We observed drug-dependent, analog control of tumor-induced cytokine secretion (Figure 1D) and cytotoxicity (Figure 1E), demonstrating that CAR expression levels within the dynamic range obtained in this model system enable tuning of CAR-T cell function.

**Figure 1:**
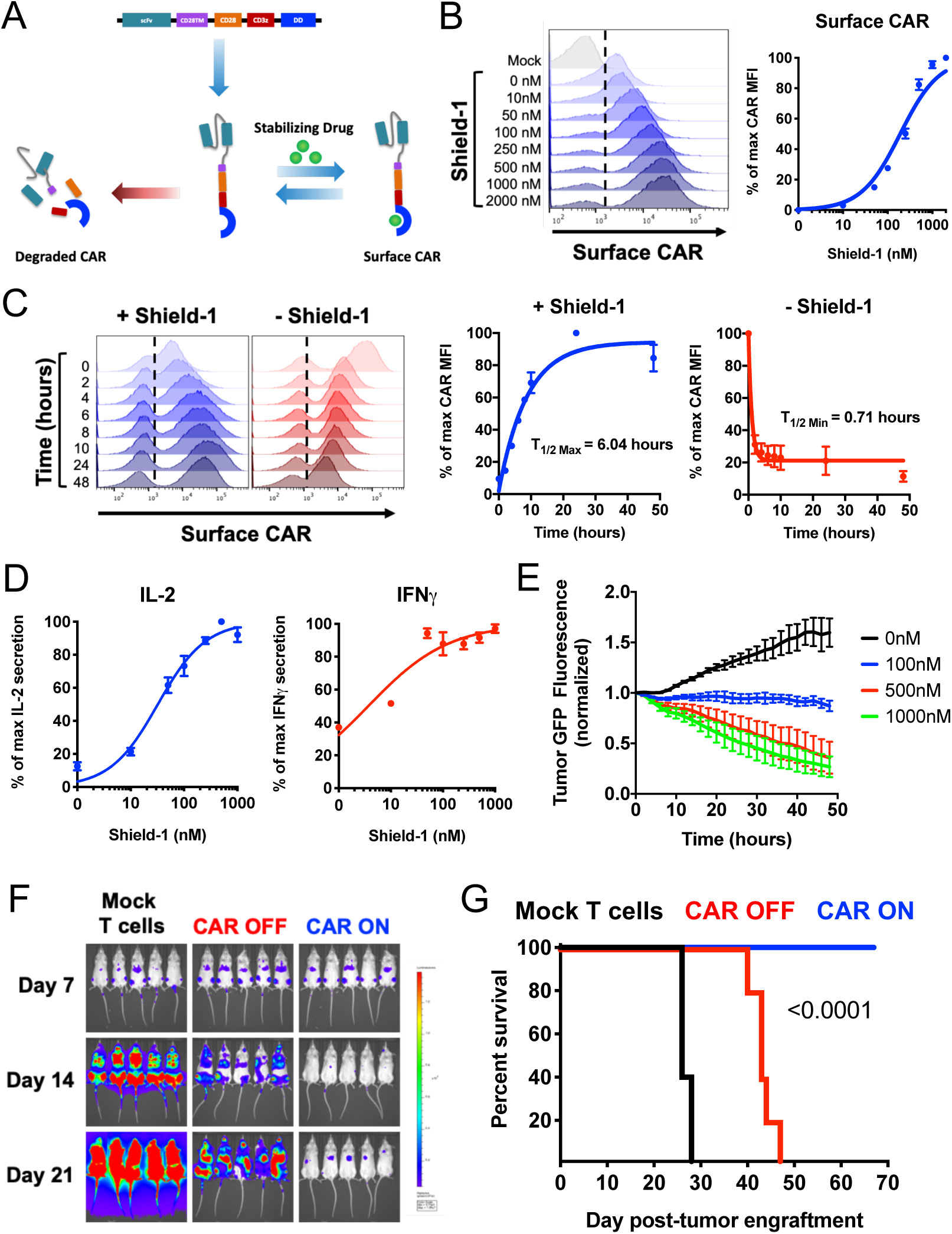
A GD2-targeting CAR modified with a destabilizing domain (DD) exhibits drug-dependent, analog control of expression and function *in vitro* and *in vivo*. **A)** Schematic depicting drug-dependent control of DD-CAR fusion protein. **B)** Flow cytometry demonstrating dose-dependent control of GD2.28*ζ*.FKBP expression via shield-1. Histogram data shows one representative donor and the dose-response curve shows mean ± SEM from 3 independent donors. **C)** Flow cytometry demonstrating ON/OFF kinetics of the GD2.28*ζ*.FKBP CAR, which was measured by adding or removing shield-1 from cell cultures and assessing CAR surface expression at the times indicated. Histograms show data from one representative donor and CAR induction and CAR disappearance curves show mean ± SEM of 3 individual donors. **D-E)** Concentrations of shield-1 were added to cells 16 hours prior to co-culture with Nalm6-GD2 leukemia. **(D)** Dose-dependent regulation of IL-2 and IFN*γ* secretion via shield-1. Error bars represent mean ± SEM of 3 individual donors. **(E)** Dose-dependent regulation of cell killing against Nalm6-GD2 leukemia via shield-1 (1:2 E:T, normalized to t=0). Error bars represent mean ± SD of triplicate wells. Representative donor from 3 individual donors. **F-G)** 1×10^6^ Nalm6-GD2 leukemia cells were engrafted in mice at day 0 and 2×10^6^ GD2.28*ζ*.ecDHFR cells expanded in the absence of TMP for 15 days *in vitro* were infused IV on day 7 post-engraftment. Mice were dosed 6 days per week with vehicle (ddH2O, CAR OFF) or 200mg/kg TMP (CAR ON). **(F)** Bioluminescence imaging of the tumor and **(G)** mouse survival data indicate TMP-dependent anti-tumor activity in CAR ON mice compared to CAR OFF mice. Representative experiment from 3 independent experiments (n=5 mice/group). Statistics were calculated using log-rank Mantel-Cox test (p<0.0001).

To interrogate DD-CAR functionality *in vivo*, we substituted the FKBP DD for an *E. coli*-derived dihydrofolate reductase DD (ecDHFR) (Iwamoto et al., 2010). Expression of GD2.28*ζ*.ecDHFR CAR is regulated by the well-tolerated, FDA-approved antibiotic, trimethoprim (TMP), and demonstrated a dynamic range comparable to that of the GD2.28*ζ*.FKBP CAR (Figure S1C). Using an NSG mouse model of Nalm6-GD2 leukemia, intraperitoneal administration of TMP induced surface CAR expression (CAR ON), upregulation of the T cell activation marker CD69 in CAR-T cells isolated from blood and spleen (Figure S1B), and robust anti-tumor potency compared to mice receiving vehicle (CAR OFF) or mock T cells (Figures 1F-G and S1C). Together, these data demonstrate drug-dependent control of DD-CAR expression and anti-tumor efficacy both *in vitro* and *in vivo*.

### Cessation of tonic CAR signaling augments CAR-T cell potency and redirects CAR-T cell fate away from exhaustion and towards a memory-like state

We next sought to determine whether drug-dependent control of DD-CAR expression could prevent exhaustion induced by tonic CAR signaling. Similar to stably expressing GD2.28*ζ* CAR-T cells (Long et al., 2015), GD2.28*ζ*.FKBP CAR-T cells cultured in the presence of shield-1 (CAR ON) exhibited antigen-independent phosphorylation of CAR CD3*ζ* (CD3*ζ*) (Figure 2A) and features of T cell exhaustion, including elevated expression of inhibitory receptors PD-1, TIM-3, and LAG-3 (Figure 2B), whereas CAR-T cells cultured in the absence of shield-1 (CAR OFF) did not. GD2.28*ζ*.FKBP or GD2.28*ζ*.ecDHFR T cells that were expanded in the CAR OFF state, but were provided drug just prior to antigen challenge (OFF/ON) exhibited superior cytotoxicity (Figure 2C), tumor-induced cytokine secretion (Figure 2D), and enhanced efficacy and cell expansion *in vivo* (Figures 2E-G). Collectively, these observations confirm the deleterious effects on T cell function caused by tonic CAR signaling and demonstrate that prevention of tonic signaling during *ex vivo* expansion augments CAR-T cell functionality.

**Figure 2:**
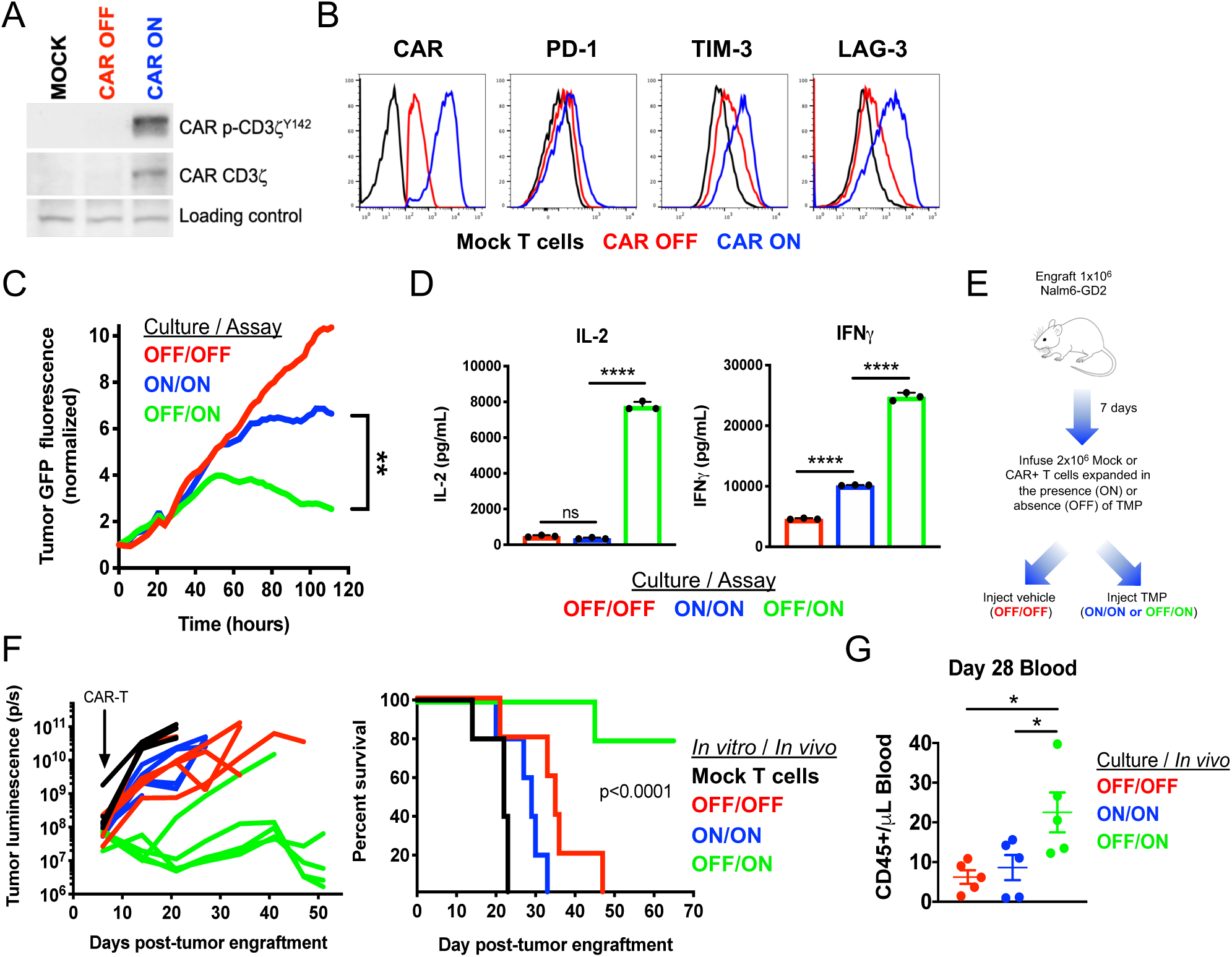
Prevention of tonic CAR signaling augments anti-tumor functionality *in vitro* and *in vivo*. **A)** Western blot demonstrating reduced total and phosphorylated CAR CD3*ζ* in GD2.28*ζ*.FKBP cells cultured in the absence of shield-1 (CAR OFF) compared to those cultured in the presence of shield-1 (CAR ON), indicating reduced tonic signaling. Representative donor from 3 individual donors. **B)** Flow cytometry demonstrates diminished inhibitory receptor expression in CAR OFF GD2.28*ζ*.FKBP CAR-T cells compared to CAR ON CAR-T cells. **C-D)** GD2.28*ζ*.FKBP CAR-T cells expanded in the absence of shield-1 for 15 days and treated with shield-1 during a co-culture assay (OFF/ON, green) exhibited **(C)** superior cytotoxicity against 143B osteosarcoma (1:8 E:T, normalized to t=0) and **(D)** increased IL-2 and IFNy secretion in response to Nalm6-GD2 leukemia compared to those treated with shield-1 during both expansion and co-culture (ON/ON, blue) and those treated with vehicle for during both expansion and co-culture (OFF/OFF, red). Data represent the mean of triplicate wells (C) or the mean ± SD. Representative donor from 3 or more donors. **E-G)** 1×10^6^ Nalm6-GD2 leukemia cells were infused IV into NSG mice at day 0 and 2×10^6^ GD2.28*ζ*.ecDHFR CAR-T cells expanded in the presence or absence of trimethoprim (TMP) for 15 days *in vitro* were infused IV on day 7 post-engraftment. Mice were dosed 6 days per week with vehicle (water, OFF/OFF) or 200mg/kg TMP (ON/ON and OFF/ON). **(F)** Tumor growth was measured via bioluminescent imaging. Mock T cell-treated or vehicle-treated mice (OFF/OFF) demonstrated rapid tumor growth (left). TMP-treated mice that received T cells grown in the absence of TMP (OFF/ON) exhibited superior tumor control and (right) prolonged survival (p<0.0001 log-rank Mantel-Cox test) compared to those that received CAR-T cells expanded in the presence of TMP (ON/ON). Representative experiment from 3 independent experiments (n=5 mice per group). **(G)** Peripheral blood was sampled on day 28 post-engraftment during the representative experiment shown in Figure 2F. Data demonstrate enhanced CAR-T cell expansion of OFF/ON CAR-T cells compared to ON/ON CAR-T cells (n=5 mice/group from 1 experiment). Statistics were calculated using two-tailed unpaired student’s t test (C, last timepoint), one-way ANOVA and Tukey’s multiple comparisons test (D,G), or log-rank Mantel-Cox test (F). *, p<0.05; **, p<0.01; ****p<0.0001; ns, p>0.05

To further explore the impact of cessation of CAR signaling on the development of exhaustion, we regulated CAR expression in HA.28*ζ*-CAR-T cells (Fig. S2A), which manifest extremely potent tonic signaling and acquire functional, transcriptomic and epigenetic hallmarks of exhaustion by D11 (Fig. 3A-C) (Lynn et al., 2019). When compared to HA.28*ζ*.FKBP CAR-T cells that were continuously cultured with shield-1 (Always ON), cells from which shield-1 was removed on D7 (Rested_D7-11_) demonstrated decreased inhibitory receptor expression and superior functionality upon tumor challenge on D11 (Figures 3A-C). To more extensively define changes in cell phenotype that occur during transition to exhaustion between D7 and D11 and the impact of rest on this process, we performed single cell analysis of 27 proteins associated with T cell exhaustion, activation, or memory (Figure S2B). Always ON cells manifested time-dependent increases in exhaustion score (normalized mean expression of PD-1, TIM-3, LAG-3, CTLA-4, BTLA, 2B4, and CD39), whereas Rested_D7-11_ cells demonstrated time-dependent increases in memory score (normalized mean expression of CD45RA, IL-7R, CD27, CD197), indicating a divergence in cell phenotypes.

**Figure 3:**
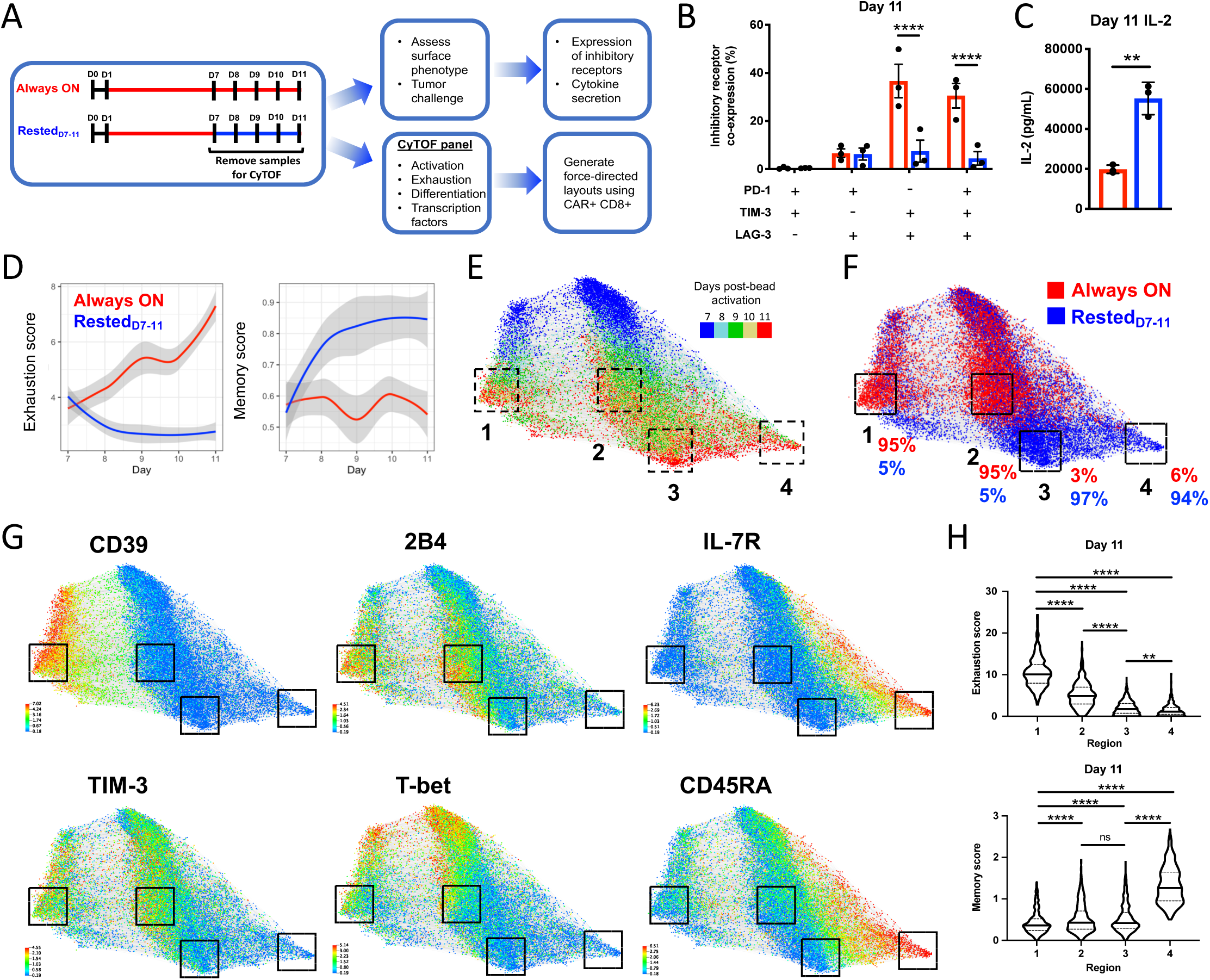
Cessation of tonic CAR signaling redirects CAR-T cell fate. **A)** Human T cells expressing the tonically signaling, high-affinity GD2-targeting CAR modified with an FKBP12 DD (HA.28*ζ*.FKBP) were cultured in the presence of shield-1 from D1-7, after which shield-1 treatment was sustained (Always ON) or discontinued (Rested_D7-11_). Cells were collected on D11 for flow cytometry and tumor co-culture assays, and on each day between D7-11 for mass cytometric analysis. **B)** Flow cytometry on D11 demonstrates reduced inhibitory receptor co-expression on Rested_D7-11_ compared to Always ON. Error bars represent mean ± SEM of 3 individual donors. **C)** IL-2 secretion in response to Nalm6-GD2 leukemia on D11. Shield-1 was added to Rested_D7-11_ 16 hours prior to tumor challenge in order to normalize CAR expression for the assay. Error bars represent mean ± SD of triplicate wells from one representative donor (n=5 individual donors). **D-H)** A force-directed layout (FDL) was constructed from 20,000 CAR+/CD8+ events analyzed by mass cytometry. Events were equally sampled from Always ON and Rested_D7-11_ cells collected on each day between D7-11 (i.e. 2,000 cells per condition/timepoint). Cells repel based on expression of 20 surface and 2 intracellular markers (Figure S2B) and grey edges connect cells from adjacent days (1 representative donor from 3 individual donors). **D)** Exhaustion score (left) for each cell was calculated as mean expression of PD-1, TIM-3, LAG-3, 2B4, CTLA-4, BTLA, and CD39 normalized to mean expression of these markers in control HA.28z.FKBP T cells cultured in the absence of shield-1. Similarly, memory score (right) was calculated using CD45RA, IL-7R, CD27, CD197 markers. Local polynomial regression fitting exhaustion and memory scores in 2,000 sampled cells from Always ON and Rested_D7-11_ conditions over time. Shaded regions indicate 95% confidence intervals. **E)** Timepoint analysis demonstrates that D11 cells are concentrated phenotypically distinct regions 1-4 (Figure S2C). **F)** Regions 1 and 2 comprise 95% D11 Always ON cells, whereas regions 3 and 4 contain 97% and 94% D11 Rested_D7-11_ cells. **G)** Marker expression demonstrates that regions 1 and 2 overexpress CD39, 2B4, TIM-3 and T-bet, whereas regions 3 and 4 demonstrate low levels of these markers and region 4 expresses high levels of IL7R and CD45RA. **H)** Violin plots of D11 cell exhaustion and memory scores contained within regions 1-4 show quartiles with a band at the mean. Statistics were calculated using one-way ANOVA and Dunnett’s multiple comparisons test. **, p<0.01; ****, p<0.0001; ns, p>0.05

Unbiased multidimensional analyses using force-directed layouts (FDLs) (Good et al., 2019; Samusik et al., 2016), which map phenotypically similar cells closely together and dissimilar cells farther apart, illustrated phenotypic trajectories towards exhaustion or memory-like cell fates. Always ON and Rested_D7-11_ CAR-T cells displayed substantial phenotypic evolution during D7-11 (Figure 3E), but manifested disparate phenotypes by D11 that were spatially distributed between 4 distinct regions on the FDL (Figure S2C). The majority of Always ON cells resided in Regions 1 or 2, while the majority of Rested_D7-11_ cells resided in Regions 3 or 4 (Figure 3F). FDLs also revealed heterogeneity within the exhausted and memory subsets, since Region 1 was defined by a higher exhaustion score and increased CD39 compared to Region 2 (Figure S2D), whereas Region 4 was defined by a higher memory score and increased expression of IL7RA and CD45RA compared to Region 3 (Figures 3G, 3H and S2E). Rested_D7-11_ cells also exhibited a sharp decline in T-bet and Blimp-1 expression levels and co-expression frequency (Figure S2F), consistent with transcriptional reprogramming of the population (Joshi et al., 2007; Rutishauser et al., 2009).

The redirection of cell fate induced by rest during D7-11 was most likely due to population-wide changes rather than outgrowth of a rare subpopulation. This is supported by the detection of modest differences in cell fold change, Ki-67 expression, and cleaved PARP (cPARP) between Always ON and Rested_D7-11_ cells (Figure S2G and S2H) (Joshi et al., 2007; Rutishauser et al., 2009). Moreover, TCF1 expression, which is associated with a progenitor exhausted cell population that retains anti-tumor functionality and is responsive to checkpoint blockade (Chen et al., 2019c; Im et al., 2016; Jadhav et al., 2019; Utzschneider et al., 2016), was similar in both Always ON and Rested_D7-11_ conditions and represented approximately 10% of total CD8+ CAR-T cells on D11 (Figure S2I). Collectively, these observations indicate that transient cessation of CAR signaling prior to exhaustion onset alters the differentiation trajectory of large fraction of CAR-T cell populations rather than inducing outgrowth of a minor subset of highly proliferative, apoptosis-resistant, or TCF1+ progenitor exhausted Rested_D7-11_ cells.

### Transient rest reverses phenotypic and transcriptomic hallmarks of exhaustion

Since cessation of tonic CAR signaling alters CAR-T cell fate during the transition to exhaustion, we hypothesized that this approach could also reprogram T cell populations on which exhaustion is already imprinted. To test this, we compared D15 Always ON HA.28*ζ*.FKBP CAR-T cells to those which had been rested from D11-15 (Rested_D11-15_, Figure 4A). Since PD-1 and PD-L1 are both expressed on activated T cells (Latchman et al., 2004) (Figure S2E), in some experiments we included a condition wherein Always ON T cells were cultured with *α*PD-1-blocking antibody from D7-15 to compare the degree of reprogramming induced by checkpoint blockade with that of transient rest. As expected, Always ON cells analyzed on D15 exhibited robust tonic CAR signaling, elevated surface expression of immune checkpoint receptors (PD-1, TIM-3 and LAG-3), and an effector-like phenotype (Figures 4B-D, S3A-C) compared to Always OFF cells. In contrast, Rested_D11-15_ cells displayed diminished tonic CAR signaling, marked reduction in checkpoint receptor expression, and increased frequencies of a stem cell memory-like subset (CD45RO-, CCR7+) compared to Always ON and Always ON + *α*PD-1 T cells. Sequential RNA sequencing on D7, D11 and D15 demonstrated that transcripts which were positively or negatively associated with exhaustion underwent rapid and complete reversal to baseline Always OFF levels in rested cells by D11 or D15, whereas inclusion of *α*PD-1 in Always ON T cell cultures minimally affected gene expression (Figures 4E and S3D).

**Figure 4:**
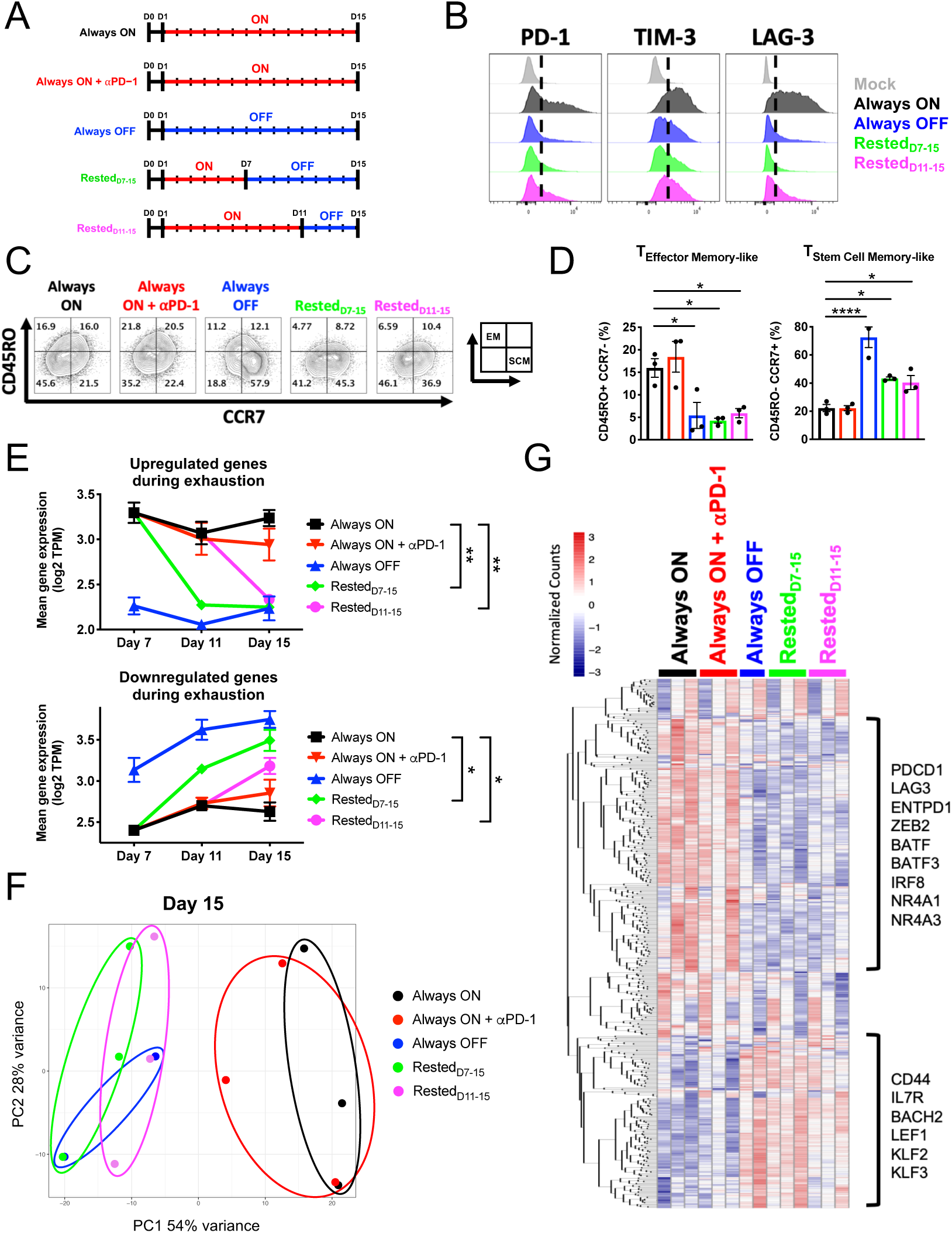
Transient rest reverses phenotypic and transcriptomic hallmarks of exhaustion. **A)** Human T cells expressing HA.28*ζ*.FKBP were cultured in the presence (Always ON, Always ON + *α*PD-1) or absence (Always OFF) of stabilizing drug (shield-1 or trimethoprim, TMP) from D1-D15, or cultured with shield-1 starting at D1 and rested for 4 or 8 days (Rested_D7-15_, Rested_D11-15_). Cells were collected on D7, D11, and D15 for FACS and bulk RNA sequencing. **B-D)** Flow cytometry of CD8+ CAR+ T cells demonstrated **(B)** reversal of inhibitory receptor expression and **(C-D)** induction of a stem cell memory-like phenotype in rested conditions compared to Always ON cells. Histograms and contour plots are representative of at least 3 donors. Bar graphs show mean ± SEM of 3 independent donors. The inset in subfigure 4C illustrates quadrants that represent effector-memory-like phenotype (EM, top-left) and stem cell memory-like phenotype (SCM, bottom-right). **E-G)** Bulk RNA sequencing analyses of mixed CD4+/CD8+ HA.28*ζ*.FKBP CAR-T cells demonstrate **(E)** rapid and dramatic alterations in exhaustion signature (Figure S3D) gene expression subsequent to rest. Error bars represent the mean ± SEM of 2-3 independent donors. Two-way ANOVA demonstrates significant differences in mean exhaustion signature gene expression between Always ON and rested conditions on D15. **(F)** Unbiased principal component analysis of D15 cells shows global transcriptomic reprogramming of rested cells to resemble Always OFF controls. **(G)** A heatmap of the top 500 genes driving principal component 1 (PC1) highlights clusters of exhaustion-and memory-associated genes that change in rested cells. Statistics were calculated using one-or two-way ANOVA and Dunnett’s multiple comparisons test. *, p<0.05; **, p<0.01; ****, p<0.0001.

Unbiased principal component analysis (PCA) at each timepoint showed significant overlap between Always OFF and rested conditions, which separated from Always ON and Always ON + *α*PD-1 cells along PC1 (Figures 4F and S3E), illustrating the high degree of transcriptomic reprogramming induced by rest. Hierarchical clustering of the top 500 genes driving the variance in PC1 identified exhaustion-associated genes (*PDCD1, ENTPD1, BATF3, NR4A1*) and memory-associated genes (*IL7R, LEF1, KLF2, BACH2)*, which were differentially expressed in exhausted cell conditions versus rested conditions (Figure 4G). Statistical analyses of D15 samples confirmed transcriptomic reversal, since Rested_D11-15_ T cells significantly upregulated memory/quiescence-associated genes (*SELL, LEF1, FOXO3*) and downregulated canonical exhaustion-associated genes (*CTLA-4, IRF4, BATF3, TOX2, NR4A3*) compared to Always ON T cells (Figure S3F). GSEA analyses confirmed that genes which were up-and down-regulated in rested conditions compared to Always ON overlap with memory and exhaustion genes, respectively, from published chronic LCMV murine studies (Wherry et al., 2007; Wherry et al., 2004) (Figure S3G). Finally, both exhausted and rested populations on D15 contained ∼1,000-3,000 unique TCR clonotypes and exhibited similar clonotypic diversity (Bolotin et al., 2017) (Figure S3H and S3I). These data are consistent with a model wherein rest-induced changes occur broadly within the population under study, rather than preferential expansion of a small subset of cells, which would result in reduced clonotypic diversity.

### Transient rest reinvigorates exhausted CAR-T cells and improves therapeutic efficacy

We hypothesized that the dramatic phenotypic and transcriptomic reprogramming of exhausted CAR-T cells induced by 4-8 days of rest would confer enhanced functionality when challenged with GD2-bearing tumor cells. Prior to antigen challenge, we treated CAR-T cells with shield-1 for approximately 16 hours to normalize CAR surface expression across experimental groups (Figure 5A). On D15, Rested_7-15_ and Rested_11-15_ T cells demonstrated a remarkable enhancement in cytotoxicity and bulk cytokine secretion compared to exhausted Always ON cells (Figure 5B, 5C and S4A). Single cell functional analyses via flow cytometry revealed that approximately 60% of Always OFF and rested CAR-T cells were polyfunctional and capable of secreting at least two cytokines, whereas less than 20% of Always ON T cell exhibited these features (Figure 5D, 5E and S4B). These data demonstrate that functional reinvigoration can be accomplished in a substantial fraction of cells following transient cessation of CAR signaling. In contrast, PD-1 blockade enhanced cytotoxicity but did not significantly increase the bulk IL-2 or IFN*γ* secretion, frequency of cytokine-secreting cells, or sensitivity to low antigen compared to rested conditions (Figures 5B, 5C, 5D, 5E, 5F, 5G, S4A and S4B), indicating that functional reinvigoration via cessation of chronic signaling is mechanistically distinct from that of checkpoint blockade. Finally, we tested whether rested HA.28*ζ*.ecDHFR CAR-T cells would exhibit more durable and potent anti-tumor efficacy *in vivo* (Figures 5H and S4D). In contrast to exhausted Always ON T cells, which failed to control tumor growth, Rested_D7-15_, and Rested_D11-15_ T cells either cured or maintained the tumor at levels comparable to Always OFF (Figure 5I, Figure S4E). These data demonstrate that transient cessation of CAR signaling substantially augments the capacity for CAR-T cells to mount a durable anti-tumor response *in vivo*.

**Figure 5:**
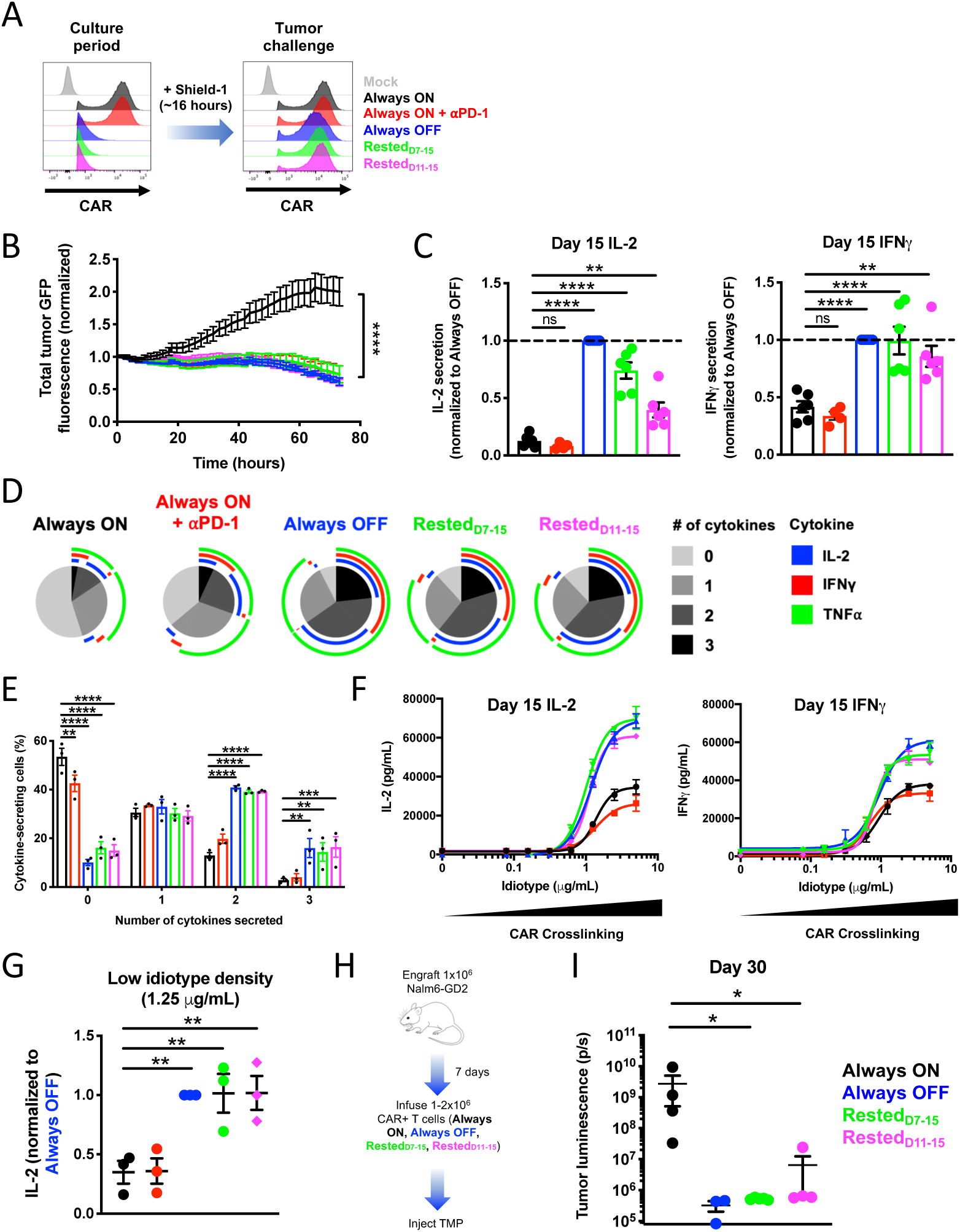
Transient rest reinvigorates exhausted CAR-T cells and improves anti-tumor function. **A)** CAR expression was normalized across all HA.28z.FKBP-expressing cells (Figure 4A) by adding shield-1 to cultures approximately 16 hours prior to *in vitro* tumor challenge. **B)** D15 incucyte assay shows that transient rest and *α*PD-1 enhances cytotoxicity in response to 143B-GL osteosarcoma (1:8 E:T, normalized to t=0) compared to exhausted Always ON controls. Error bars represent mean ± SD of 3 triplicate wells from one representative donor (n=3 individual donors). **C)** D15 co-culture assay with Nalm6-GD2 leukemia demonstrates that transient rest, but not *α*PD-1, augments IL-2 and IFN*γ* secretion. Error bars represent mean ± SEM of 4-6 individual donors. **D-E)** Intracellular cytokine staining and flow cytometry analyses of CD8+ CAR+ T cells demonstrate that rest reduces the frequency of non-responsive cells and increases the frequency of polyfunctional cells. SPICE analysis from 1 representative donor was conducted in (D). Error bars in (E) represent mean ± SEM of 3 individual donors. **F-G)** CAR-T cells were crosslinked with immobilized 1A7 anti-CAR idiotype antibody for 24 hours. **(F)** Non-linear dose-response curves demonstrate that T cell rest, but not PD-1 blockade, augments IL-2 and IFN*γ* secretion in response to both low and high idiotype densities. Error bars represent mean ± SD of 3 triplicate wells from one representative donor (n=3 individual donors). **(G)** IL-2 secretion in response to low density (1.25*μ*g/mL) anti-CAR idiotype was normalized to secretion levels from Always OFF cells. Error bars represent mean ± SEM of 3 individual donors. **H-I)** 1×10^6^ Nalm6-GD2 leukemia cells were engrafted in mice on day 0 and 1-2×10^6^ HA.28*ζ*.ecDHFR CAR-T cells expanded for 15 days *in vitro* (as depicted in Figure 4A) and were infused IV on day 7 post-engraftment. Mice were dosed with 200mg/kg TMP 6 days/week. **(I)** Bioluminescent imaging on day 30 post-engraftment demonstrates augmented control of tumor growth in Always OFF and rested conditions compared to Always ON cells. Error bars represent mean ± SEM of 3-5 mice from 1 representative experiment (n=3 independent experiments). Statistics were calculated using one or two-way ANOVA and Dunnett’s multiple comparisons test. *, p<0.05; **, p<0.01; ***, p<0.001; ****, p<0.0001

### Rest induces wholescale remodeling of the exhaustion-associated epigenome

To determine the impact of transient cessation of CAR signaling on the epigenome, we used ATAC-seq (Buenrostro et al., 2013) to analyze differences in chromatin accessibility between Always ON, Always OFF, Rested_7-15_ and Rested_11-15_ HA.28*ζ*.FKBP CD8+ CAR-T cells (Figure 6A). Temporal analysis of epigenetic reprogramming in this model system revealed that T cells that experienced continuous tonic CAR signaling (Always ON) underwent the most dramatic alterations in chromatin accessibility within the first 7 days (∼48,000 peak changes), with fewer changes occurring between D7-11 (∼2,300 peak changes) and D11-15 (∼2,000 peak changes) (Figures 6B and S5A). Remarkably, 4 days of rest occurring between D7-11 (Rested_D7-11_) or D11-15 (Rested_D11-15_) was associated with ∼19,000 and ∼15,500 peak changes, respectively (Figure 6B), many of which were differentially accessible compared to Always ON cells (Figure S5B). These included exhaustion-associated genes *ENTP1* and *BATF3* and the stemness-associated gene *TCF7* (Figures 6C). These results align with our observations of rapid phenotypic, functional and transcriptional reprogramming following 4 days of rest.

**Figure 6:**
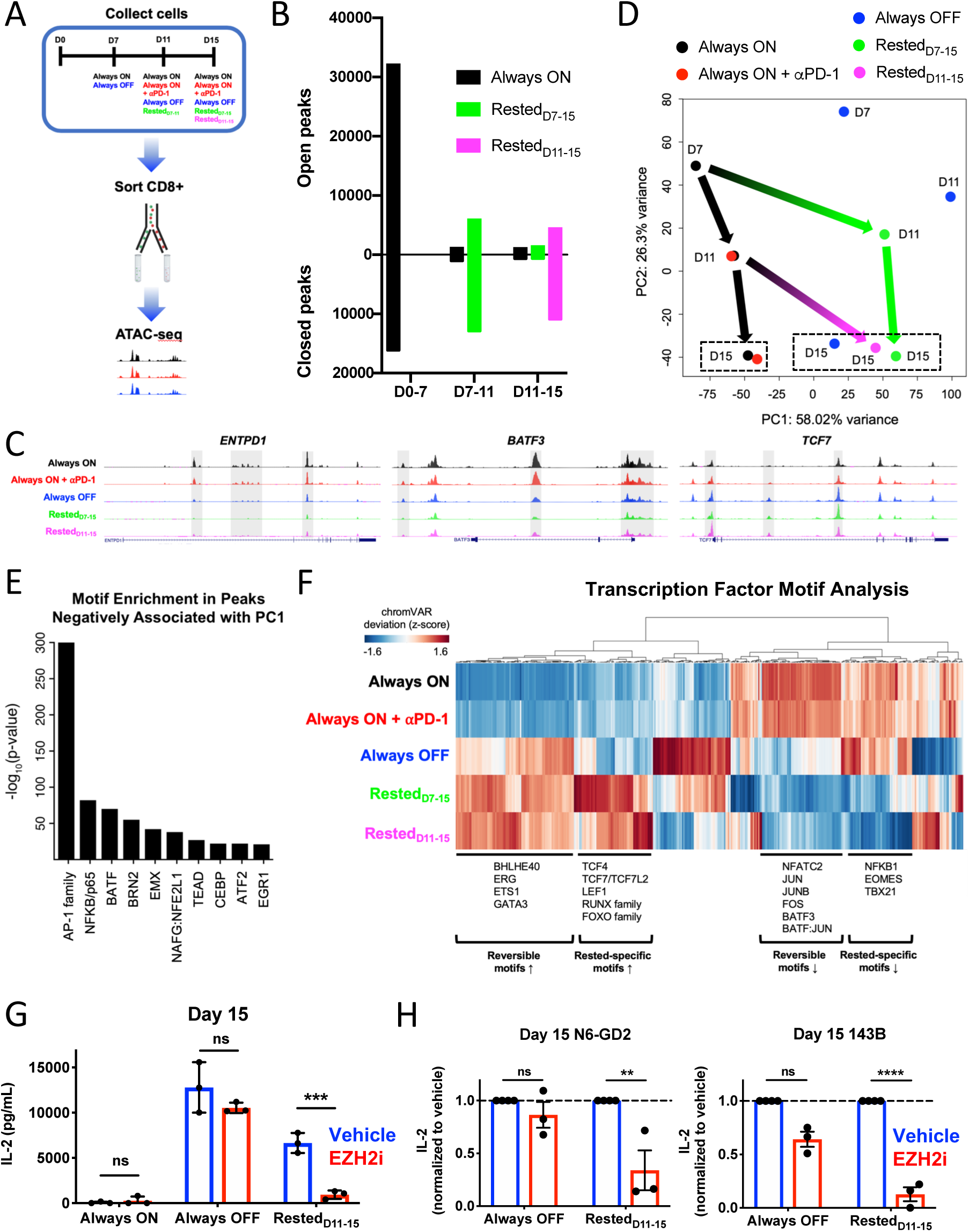
Rested CAR-T cells exhibit wholescale remodeling of the exhaustion-associated epigenome. **A-F)** HA.28ζ.FKBP CAR-T cells were cultured as depicted in Fig. 4A, and **(A)** CD8+ T cells were sorted on D0 (prior to bead activation), D7, D11, and D15 for ATAC-seq. **B)** Peak accessibility changes between timepoints was calculated based on p adjusted<0.05. Bar graphs display merged data from 2-3 donors. **C)** Differentially accessible regions in the *ENTPD1*, *BATF3*, and *TCF7* loci on D15. Representative donor of 2-3 donors. **D)** Unbiased principal component analysis of chromatin accessibility was assessed across all timepoints. Always ON and Always ON + αPD-1 are separated from Always OFF, Rested_D7-15_, and Rested_D11-15_ along PC1 (58% variance). Green and magenta arrows indicate global epigenetic remodeling upon rest. Dashed boxes indicate D15 samples. Each dot represents merged data from 2-3 individual donors. **E)** TF motifs enriched in the top 5000 peaks that are negatively associated with PC1 indicate a strong AP-1 family signature in exhausted Always ON and Always ON + αPD-1 samples that is reversed following T cell rest. **F)** Hierarchical clustering of differentially accessible TF motifs in D15 samples identified reversible and rested-specific clusters of TF motifs. The heatmap was generated using merged data from 2-3 individual donors. **G-H)** Always ON, Always OFF, and Rested_D11-15_ CAR-T cells were cultured with 1μM tazemetostat (EZH2i) or vehicle from D11-15. Shield-1 was added to CAR-T cells approximately 16 hours prior to co-culture with N6-GD2 leukemia or 143B-GL osteosarcoma, and EZH2i was washed out immediately prior to co-culture. **G)** IL-2 secretion in response to N6-GD2 showing mean ± SD of triplicate wells from 1 representative donor (n=3 individual donors). **(H)** IL-2 secretion in response to N6-GD2 or 143B-GL was normalized to vehicle-treated controls, where error bars represent mean ± SEM of 3 individual donors. Statistics were calculated using two-way ANOVA and Dunnett’s multiple comparisons test. **, p<0.01; ***, p<0.001; ns, p>0.05

Unbiased Pearson correlation and PCA analyses indicate that rest results in global, wholescale remodeling of the epigenome. Always ON cells treated with or without anti-PD-1 cluster together along PC1 on D11 (66.51% variance) and D15 (58% variance) and exhibit highly correlated genomic accessibility profiles (Figures 6D, S5C and S5D). In contrast, Rested CAR-T cells exhibit clear separation from Always ON and cluster together with Always OFF cells along PC1 (Figure 6D, S5C and S5D).

AP-1 family TF motifs (BATF, BATF3, JUNB, NFATC2), which are known to promote T cell exhaustion (Kurachi et al., 2014; Lynn et al., 2019; Man et al., 2017; Martinez et al., 2015; Scott-Browne et al., 2016), were highly enriched in accessible regions of the Always ON epigenome, but rendered less accessible following rest (Figures 6E and 6F). TF binding motifs for genes that have been implicated in T cell memory were exclusively accessible in rested conditions (TCF7/TCF7L2, LEF1, RUNX family, FOXO family) (Hu and Chen, 2013; Jadhav et al., 2019; Kratchmarov et al., 2018; Michelini et al., 2013; Milner et al., 2017; Rao et al., 2012; Sullivan et al., 2012; Wang et al., 2018; Xing et al., 2016), whereas those associated with T cell exhaustion (EOMES, TBX21) demonstrated exaggerated inaccessibility compared to Always OFF control cells (Figure 6F), raising the prospect that T cell rest could induce a unique program to drive the development of memory-like T cells. Gene set enrichment of the genes associated with differentially accessible peaks at D15 revealed biological processes that are enriched (packaging of telomeres, G1 phase) or diminished (Akt signaling, apoptosis, negative regulation of Wnt signaling, chromatin silencing complex) in rested conditions (Figure S5E), consistent with induction of a quiescent/memory T cell phenotype following rest.

Recent studies suggest that the epigenetic modifier, enhancer of zeste homolog 2 (EZH2), which catalyzes trimethylation on histone 3 at lysine 27 (H3K27me3) as a subunit of the polycomb repressive complex 2 (PRC2), prevents hematopoietic stem cell exhaustion and is critical for T cell differentiation, maintenance of T cell memory, and antitumor immunity (Chen et al., 2019b; He et al., 2017; Kamminga et al., 2006). To test whether EZH2 function is involved in reversal of exhaustion in this model, we rested Always ON exhausted CAR-T cells in the presence of the EZH2 selective inhibitor tazemetostat (EZH2i) from D11-15. EZH2i treatment had no effect on Always ON or Always OFF T cells, but attenuated the rescued IL-2 secretion in Rested_D11-15_ CAR-T cells compared to vehicle controls, suggesting that rest induces active chromatin remodeling in exhausted CAR-T cells via EZH2 (Figures 6G and 6H). Of note, EZH2 inhibition did not affect rest-induced reversal of the exhausted cell surface phenotype (Figure S5F), consistent with the observation that the epigenetic mechanisms governing exhausted T cell phenotype and dysfunction are distinct (Scott et al., 2019).

### Reinvigoration of exhausted CAR-T cells using the Src kinase inhibitor dasatinib

We and others recently demonstrated that dasatinib, an FDA-approved tyrosine kinase inhibitor, suppresses CAR-T cell activation via rapid and reversible antagonism of proximal T cell receptor (TCR) signaling kinases (Mestermann et al., 2019; Weber et al., 2019). Consistent with our previous findings, dasatinib-treated HA.28*ζ* CAR-T cells exhibited undetectable phosphorylation of CAR CD3*ζ* and ERK1/2 compared to those treated with vehicle (Figure S6A), indicating that dasatinib potently suppresses tonic signaling in CAR-T cells. We hypothesized that dasatinib-induced inhibition of tonic CAR signaling could be used to induce rest and reverse exhaustion. Indeed, Dasatinib_D4-15_, Dasatinib_D7-15_ and Dasatinib_D11-15_ cells exhibited diminished inhibitory receptor expression, functional reinvigoration as measured by antigen-induced cytokine production (Figures S6B and S6C), and improved control of tumor growth following adoptive transfer (Figures 7A). CAR-T cells expressing GD2.BB*ζ*, which also exhibit tonic signaling (Long et al., 2015), were also enhanced following *ex vivo* dasatinib treatment in a 143B osteosarcoma xenograft model (Figure S6D), indicating that dasatinib provides an approach to mitigate deleterious CAR signaling in pre-clinical or clinical CAR manufacturing settings.

**Figure 7:**
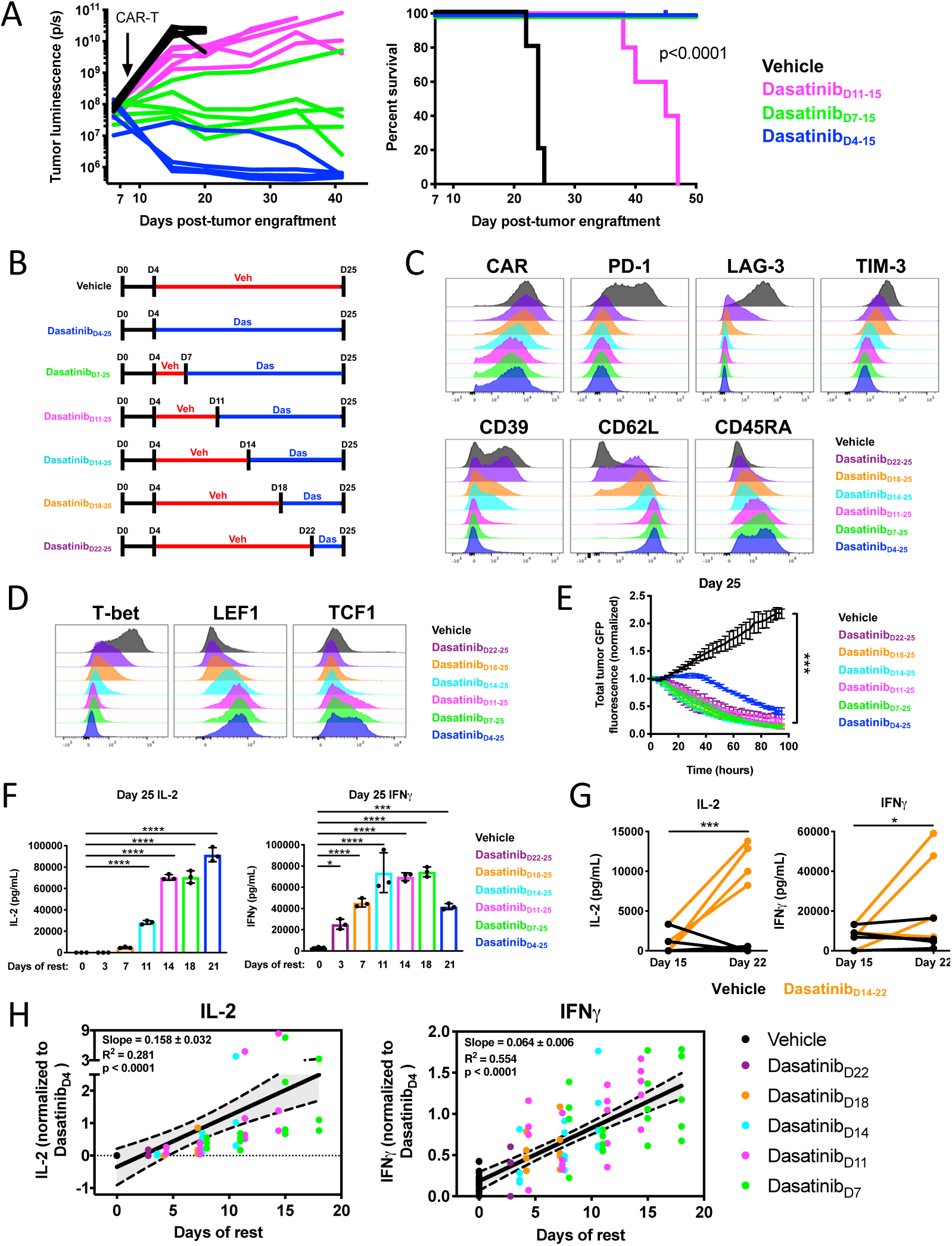
Reinvigoration of exhausted CAR-T cells using the Src kinase inhibitor dasatinib. **a)** 1×10^6^ Nalm6-GD2 leukemia cells were engrafted in mice on day 0 and 1×10^6^ D15 HA.28*ζ* CAR-T cells expanded with vehicle (DMSO) or dasatinib (4-11 days) *in vitro* were infused intravenously on day 7 post-engraftment. (left) Bioluminescent imaging and (right) survival curves demonstrate augmented control of tumor growth in dasatinib-treated conditions compared to vehicle-treated controls (p<0.0001 log-rank Mantel-Cox test), wherein the duration of rest via dasatinib correlates to anti-tumor potency. Representative experiment from 3 individual experiments (n=5 mice per group). **B-H)** Human CAR-T cells expressing HA.28*ζ* were cultured with vehicle (DMSO) or dasatinib for 3-21 days. Cells were collected days 15, 18, 22, and 25 for FACS and/or functional assays. Dasatinib was removed from the culture medium approximately 16 hours prior to *in vitro* tumor challenge. **C-D)** D25 FACS data demonstrates that short-or long-term dasatinib treatment results in **(C)** diminished exhaustion marker expression, increased memory marker expression, and **(D)** profound changes in expression of exhaustion-and stemness-associated transcription factors. Representative donor of 3 individual donors. **E)** D25 co-culture assay using Nalm6-GD2 leukemia (1:4 E:T, normalized to t=0) demonstrates that 3-21 days of rest via dasatinib treatment reverses defects in cytotoxicity in exhausted, vehicle-treated HA.28*ζ* CAR-T cells. Error bars represent mean ± SD of triplicate wells from 1 representative donor (n=3 individual donors). **F)** D25 IL-2 and IFN*γ* secretion indicates that rest via dasatinib augments cytokine secretion in response to Nalm6-GD2 leukemia. Error bars represent mean ± SD of triplicate wells from one representative donor (n=3 individual donors). **G)** DasatinibD14-22 cells exhibit increased IL-2 and IFN*γ* on D22 compared to D15 (prior to dasatinib treatment), indicating reversal of cytokine dysfunction. Data represents 4 individual donors. **H)** Co-culture assays with Nalm6-GD2 leukemia were conducted on D15, D18, D22, and D25. IL-2 and IFN*γ* secretion levels were normalized to DasatinibD4-25 for each individual timepoint and plotted based on the duration of rest (i.e. number of days of dasatinib treatment). Dot color represents the day on which dasatinib was added. Linear regression analyses demonstrate that the duration of rest Vehicle significantly correlates with the degree to which exhausted CAR-T cells are reinvigorated. Graphs display data from 4-5 individual donors (IL-2, n=63 data points. IFN*γ*, n=85 data points). D15 IL-2 and IFN*γ* data from one donor was not assessed. D18 IL-2 from one donor was omitted due to technical artifacts. Statistics were calculated using two-way ANOVA and Bonferroni’s multiple comparisons test. *, p<0.05; ***, p<0.001; ****, p<0.0001

These experiments also revealed, however, that Dasatinib_D11-15_ cells did not demonstrate equivalent anti-tumor functionality to Dasatinib_D4-15_ and Dasatinib_D7-15_ cells, raising the prospect that these cells encountered a point of irreversibility in this model system and/or they experienced an inadequate period of rest to restore functionality. To determine which of these factors was responsible for the incomplete reinvigoration observed in Dasatinib_D11-15_ cells, we sought to interrogate exhaustion reversibility using a more protracted *in vitro* time course, wherein cells were rested in increments of 3-4 days from D4 until D25 (Figure 7B). Since dasatinib completely suppresses CAR-T cell signaling and function (Figure S6A) (Mestermann et al., 2019; Weber et al., 2019), whereas DD-CARs in the OFF state exhibited some leakiness in expression and function (Figures 1, S1B and S1C), we opted to utilize dasatinib to induce rest in the protracted model system. All groups of dasatinib-treated HA.28*ζ* CAR-T cells analyzed on D25 demonstrated diminished exhaustion marker expression and increased stem cell memory-associated CD62L and CD45RA expression (Figures 7C and S6E). Dasatinib-treated groups also exhibited diminished expression of T-bet and increased expression of stemness-associated transcription factors LEF1 and TCF1 (Figures 7D and S6F), which corroborates D15 epigenetic changes to these factors or their binding motifs (Figures 6C and 6F).

Importantly, a more prolonged period of rest initiated on D11 (Dasatinib_D11-25_) resulted in complete functional restoration (Figure 7F), consistent with a model wherein the incomplete reinvigoration observed on D15 (Figures 7A and S6C) reflected an insufficient period of rest rather than irreversibility of the exhaustion program (Figures 7E-G). This model is further evidenced by a strong correlation observed between the degree of functional reinvigoration and duration of rest, which was independent of the time at which rest was initiated (Figures 7H, S6G and S6H). Collectively, these results demonstrate that pharmacologic antagonism of TCR signaling kinases reinvigorates exhausted CAR-T cells, raising the prospect that transient suppression of chronic TCR/CAR activation signals can reverse the exhaustion program.

## DISCUSSION

Chronic antigen stimulation induces T cell exhaustion, a differentiation state associated with a heritable epigenetic imprint that is distinct from that found in effector and memory T cells (Pauken et al., 2016; Philip et al., 2017; Schietinger et al., 2012; Schietinger et al., 2016; Sen et al., 2016; Wherry et al., 2007; Zajac et al., 1998). PD-1 blockade can reinvigorate exhausted T cells (Barber et al., 2006) and therapeutic agents targeting the PD-1/PD-L1 axis have improved outcomes for many patients with cancer, but durable anti-tumor effects occur in only a fraction of those treated (Ribas and Wolchok, 2018). One explanation for the limited efficacy of PD-1/PD-L1 blockade relates to its inability to reverse the exhaustion-associated epigenetic imprint (Pauken et al., 2016). Recent studies have identified TOX, TOX2, and AP-1 family members as central regulators of human T cell exhaustion that promote widespread transcriptional and epigenetic dysregulation (Alfei et al., 2019; Khan et al., 2019; Scott et al., 2019; Seo et al., 2019; Wang et al., 2019; Yao et al., 2019). These findings have enabled new approaches to mitigate exhaustion, including enforced expression of c-Jun (Lynn et al., 2019) or CRISPR-mediated deletion of TOX, TOX2, or NR4A family TFs (Chen et al., 2019a; Seo et al., 2019). However, such modifications do not result in wholescale remodeling of the exhaustion-associated epigenome, suggesting that exhaustion is an epigenetically fixed cell state.

In this report, we modified a validated *in vitro* model of CAR-associated T cell exhaustion, wherein tonic CAR signaling induces hallmark phenotypic, functional, transcriptomic and epigenetic features of exhaustion within 11 days (HA.28*ζ*) (Lynn et al., 2019), to enable precise, drug-dependent control of CAR signaling. Consistent with murine models of viral infection wherein antigen clearance induces T cell memory rather than exhaustion (Joshi et al., 2007; Rutishauser et al., 2009; Wherry et al., 2007; Wherry et al., 2004), early cessation of tonic CAR signaling (Rested_D7-11_) redirected T cell differentiation away from exhaustion and towards a memory-like state (Figure 3). Importantly, when inhibition of CAR signaling was delayed until D11, a time when cells had already acquired hallmark features of T cell exhaustion, we also observed impressive functional reinvigoration associated with global phenotypic, transcriptomic and epigenetic reprogramming (Figures 4-7). Similar results were obtained using the Src kinase inhibitor dasatinib to inhibit CAR signaling, wherein functional reinvigoration was observed even in CAR-T cells subjected to tonic signaling for longer periods (Figure 7).

Several groups have sought to identify subsets of exhausted T cells that retain the capacity for reversal of the exhaustion program (Im et al., 2016; Miller et al., 2019; Utzschneider et al., 2016; Utzschneider et al., 2013). In murine models of LCMV, “progenitor exhausted” T cells, which are defined by expression of the stemness transcription factor TCF1 (gene name *Tcf7*), are essential for providing the proliferative burst observed following PD-1 blockade (Chen et al., 2019c; Im et al., 2016; Jadhav et al., 2019; Miller et al., 2019; Utzschneider et al., 2016). Similarly, murine tumor models have revealed that exhaustion reprogrammability is associated with increased accessibility at the *Tcf7* locus (Philip et al., 2017). Finally, in melanoma patients treated with PD-1/PD-L1 blockade, lower levels of *Tcf7* expression served as a biomarker for resistance to this therapy (Sade-Feldman et al., 2019). One model to explain our findings posits that cessation of tonic CAR signaling induces preferential expansion of pre-dysfunctional, progenitor exhausted T cells. Consistent with this, rested CAR-T cell populations exhibited increased accessibility at the *Tcf7* locus, enriched *Tcf7* binding motifs within accessible regions of the genome, and increased frequency of TCF1+ cells compared to exhausted CAR-T cells (Figures 6F and 7D, S2I and S6F). However, TCF1+ cells in reinvigorated dasatinib-treated groups did not co-express PD-1 or other immune checkpoint receptors (Figures 6C and 6D), a canonical feature of progenitor exhausted T cells (Chen et al., 2019c; Im et al., 2016; Jadhav et al., 2019; Miller et al., 2019; Utzschneider et al., 2016).

An alternative model posits that some exhausted cell populations that have acquired the hallmark epigenetic imprint retain the capacity for epigenetic remodeling to resemble healthy, non-tonically signaling CAR-T cells. Results demonstrating that tazemetostat, an EZH2 inhibitor, prevents functional reinvigoration induced by rest in this model system are consistent with epigenetic reprogramming rather than enrichment of progenitor exhausted cells (Figure 6). Moreover, clonotypic analyses demonstrated similarly high levels of TCR diversity in both exhausted and rested cell populations, indicating that transcriptional and epigenetic alterations induced by rest are unlikely to occur via preferential expansion of a small subset of clones. Future studies are warranted to better define the specific cell populations that may undergo epigenetic reprogramming following T cell rest and to identify the precise role of EZH2 on remodeling of the exhaustion-associated epigenetic imprint.

Irrespective of mechanism, these results demonstrate that cessation of CAR signaling is a potent maneuver that results in augmented function in cell populations that are transitioning to exhaustion or in those already imprinted with hallmark features of exhaustion, and is distinct from that which can be achieved with PD-1/PD-L1 blockade. Advances in synthetic biology have enabled the development of several tunable CAR platforms which have been optimized to mitigate CAR-mediated toxicity (Bejestani et al., 2017; Cho et al., 2018; Leung et al., 2019; Rodgers et al., 2016; Wu et al., 2015). Our data suggest that such platforms, including the DD-CAR system described here, may enhance CAR-T cell potency as a result of temporal control of CAR-T cell signaling. Pulsed CAR expression instead of constitutive expression may allow for rest phases that collectively mitigate/reverse exhaustion and augment durability. Consistent with this, a recent study testing a regulatable CD19-targeting CAR against non-malignant B cells in a murine model demonstrated that CAR-T cells provided a longer rest phase exhibited superior antigen-induced expansion compared to those that received rest for a shorter period (Viaud et al., 2018). Similarly, short-term dosing of dasatinib *in vivo* augmented CD19-targeting CAR-T cell responses in a leukemia xenograft model (Mestermann et al., 2019), which we speculate may result from the phenomena described here. Furthermore, our work demonstrates that pharmacologic inhibition of tonic CAR signaling via dasatinib could be used to prevent T cell exhaustion during *ex vivo* expansion of clinical CAR-T cell products, and thereby enhance functionality following adoptive transfer (Figures 7 and S6).

The findings presented here also indicate that therapies designed to transiently inhibit TCR signaling might enhance functionality of exhausted, non-engineered T cell populations. Indeed, improved antigen-specific T cell function has been associated with antigen clearance in humans with hepatitis C infection treated with direct acting anti-viral therapies (Burchill et al., 2015; Kohli et al., 2015; Martin et al., 2014; Shrivastava et al., 2017; Wieland et al., 2017). In this setting, a sizable fraction of patients with ongoing viral replication at the time of cessation of anti-viral therapy ultimately clear the virus, consistent with reinvigoration of antiviral immunity. Similarly, an immunomodulatory effect of dasatinib on T cells has been associated with improved anti-tumor immunity (Hekim et al., 2017). Collectively, these observations suggest that transient cessation of TCR signaling could provide a widely applicable but underappreciated approach to enhance functionality in populations of exhausted human T cells.

In summary, we demonstrate that transient cessation of CAR signaling can reverse dysfunction and induce epigenetic reprogramming in exhausted human CAR-T cell populations. These results suggest that CAR-T cell therapeutics designed to incorporate periods of rest may exhibit increased potency compared to constitutive platforms, and raise the prospect that targeting proximal TCR/CAR signaling kinases may represent a novel immunotherapeutic strategy for mitigating T cell exhaustion.

## METHODS

| REAGENT OR RESOURCE | SOURCE | IDENTIFIER |
| --- | --- | --- |
| **Mass Cytometry Antibodies and Reagents** |  |  |
| Anti-cPARP (clone Asp214) | BD | 300443 |
| Anti-CD27 (clone 323) | Biologend | 302839 |
| Anti-CD62L (clone DREG-56) | Biologend | 304835 |
| Anti-CD4 (clone RPT-T4) | Fluidigm | 3145001B |
| Anti-CD8 (clone RPT-T8) | Fluidigm | 3146001B |
| Anti-CD28 (clone CD28.2) | Biologend | 302937 |
| Anti-PDL1 (clone 29E.2A3) | Fluidigm | 3148017B |
| Anti-CD45RO (clone UCHL1) | Fluidigm | 3149001B |
| Anti-OX-40 (clone ACT35) | Fluidigm | 3150023B |
| Anti-2B4 (clone C1.7) | Biologend | 329502 |
| Anti-TIM-3 (clone F38-2E2) | Fluidigm | 3153008B |
| Anti-CXCR3 (clone G025H7) | Fluidigm | 3156004B |
| Anti-4-1BB (clone 4B4-1) | Fluidigm | 3158013B |
| Anti-Blimp-1 (clone UCHT1) | Biologend | 300402 |
| Anti-T-bet (clone 4B10) | Biologend | 3160010B |
| Anti-CTLA-4 (clone 14D3) | Fluidigm | 3161004B |
| Anti-CD25 (clone M-A251) | BD | 555430 |
| Anti-BTLA (clone MIH26) | Fluidigm | 3163009B |
| Anti-CD39 (clone A1) | Biologend | 328221 |
| Anti-LAG-3 (clone 874501) | Fluidigm | 3165028B |
| Anti-CCR7 (clone G043H7) | Fluidigm | 3167009A |
| Anti-IL-7R (clone A019D5) | Fluidigm | 3168017B |
| Anti-CD45RA (clone HI100) | Fluidigm | 3170010B |
| Anti-Ki-67 | Biologend | 350523 |
| Anti-CD95 (clone DX2) | Biologend | 305631 |
| Anti-PD-1 (clone EH12.2H7) | Fluidigm | 3174020B |
| Anti-Eomes (clone WD1928) | eBioscience | 14-4877-82 |
| DNA intercalator | Fluidigm | 201191B |
| Cisplatin | Fluidigm | 201064 |
| Cell-ID 20-Plex Pd Barcoding Kit | Fluidigm | 201060 |
| EQ Four Element Calibration Beads | Fluidigm | 201078 |
| **Flow Cytometry Antibodies and Reagents** |  |  |
| Anti-14g2A idiotype antibody (clone 1A7) | NIH |  |
| Anti-CD4 (clone SK3) | BD | 563550 |
| Anti-CD8 (clone SK1) | BD | 564912 |
| Anti-PD-1 (clone eBioJ105) | eBioscience | 25-2799-42 |
| Anti-TIM-3 (clone F38-2E2) | Biologend | 345030 |
| Anti-LAG-3 (clone 3DS223H) | eBioscience | 12-2239-42 |
| Anti-CD39 (clone A1) | Biologend | 328226 |
| Anti-CD62L (clone DREG-56) | BD | 562719 |
| Anti-CCR7 (clone 150503) | BD | 562555 |
| Anti-CD45RO (clone UCHL1) | BD | 25-0457-42 |
| Anti-CD45RA (clone L48) | BD | 649458 |
| Anti-T-bet (clone 4B10) | Biolegend | 644819 |
| Anti-LEF1 (clone C12A5) | Cell Signaling | 14440S |
| Anti-TCF7 (clone C63D9) | Cell Signaling | 6709S |
| Anti-IL-2 (clone MQ1-17H12) | Biolegend | 500326 |
| Anti-IFNγ (clone 4S.B3) | Biolegend | 502530 |
| Anti-TNFα (clone MAb11) | Biolegend | 502940 |
| Anti-CD69 (FN50) | Biolegend | 310930 |
| Anti-CD107a (clone H4A3) | Biolegend | 328634 |
| Monensin | eBioscience | 00-4505-51 |
| Fixable Viability Dye | eBioscience | 65-0866-14 |
| FoxP3 Transcription Factor Staining Buffer Set | eBioscience | 00-5523-00 |
| CountBright Absolute Counting Beads | Thermo Scientific | C36950 |
| DyLight 650 Microscale Antibody Labeling Kit | Thermo Fisher | 84536 |
| DyLight 488 Microscale Antibody Labeling Kit | Thermo Fisher | 53025 |

## CONTACT FOR REAGENT AND RESOURCE SHARING

Further information and requests for resources and reagents should be directed to and will be fulfilled by the Lead Contact, Crystal L. Mackall.

## EXPERIMENTAL MODEL AND SUBJECT DETAILS

### Primary human T cells

Anonymous healthy donor buffy coats were obtained from the Stanford University Blood Center (Stanford, CA) under a University Institutional Review Board-exempt protocol. Primary CD3+ T cells were subsequently isolated from buffy coats via negative selection (described in METHOD DETAILS).

### Cell Lines

143B osteosarcoma cells (ATCC) were stably transduced to express GFP, firefly luciferase, and CD19 (143B-GL). Nalm6 B-ALL cell line that was stably transduced to express GFP and firefly luciferase (Nalm6-GL) was provided by David Barrett. Nalm6-GL was co-transduced with cDNAs for GD2 synthase and GD3 synthase in order to induce overexpression of GD2 (Nalm6-GD2). Single cell clones were then chosen for high antigen expression. All cell lines were cultured in complete media (RPMI supplemented with 10% FBS, 10mM HEPES, 2mM GlutaMAX, 100 U/mL penicillin, and 100ug/mL streptomycin (Gibco)).

### Murine xenograft models

NOD/SCID/IL2R*γ^−/−^* (NSG) mice were bred, housed, and treated under Stanford University APLAC-approved protocols. 6-8 week old male or female mice used in experiments were healthy and immunocompromised, drug and test naïve, and were not involved in previous procedures. Mice were housed in sterile cages in the barrier facility located in the Veterinary Service Center (VSC) at Stanford University with a 12-hour light/dark cycle. Mice were monitored daily by the VSC husbandry staff and were euthanized upon manifestation of hunched posture, impaired mobility, rough coat, paralysis, or tumor sizes that exceeded the limits designated in approved animal protocols. For xenograft experiments using Nalm6-GD2, 1×10^6^ tumor cells in 200uL PBS were engrafted via intravenous injection. 1-2×10^6^ CAR-T cells suspended in 200uL PBS were subsequently infused intravenously on day 7 post-engraftment. Tumor burden was monitored via bioluminescence imaging using a Spectrum IVIS instrument and quantified using Living Image software (Perkin Elmer). For xenograft experiments using 143B cells, 1×10^6^ tumor cells in 100uL PBS were engrafted via intramuscular injection into the flank and 10×10^6^ CAR-T cells suspended in 100uL PBS were infused intravenously on day 3 post-engraftment. Tumor size was monitored via caliper measurements. Mice were randomized to ensure equal tumor burden before CAR-T cell treatment. For experiments involving GD2.28*ζ*.ecDHFR or HA.28*ζ*.ecDHFR-expressing CAR-T cells, 200uL of vehicle (ddH2O, Thermofisher, # 10-977-023) or 200uL of 200mg/kg trimethoprim lactate resuspended in ddH2O (TMP, Bioworld, #42010018-3) was injected into mice intraperitoneally 6 days/week.

## METHOD DETAILS

### CAR construct design

GD2.28*ζ*, GD2.BB*ζ*, and HA.28*ζ* CARs (Long et al., 2015; Lynn et al., 2019) were constructed in pELNS lentiviral vectors and/or MSGV retroviral vectors. Codon-optimized gene fragments that included a portion of the CAR CD3z domain sequence (downstream of the BmgBI restriction site) and FKBP12 or ecDHFR destabilizing domains were generated by IDT. Fragments were then inserted into pELNS.GD2.28*ζ* and pELNS.HA.28*ζ* vectors at the 5’ BmgBI restriction cut site within the CAR CD3z domain and the 3’ SalI site downstream of the stop codon using T4 DNA Ligase (New England Biolabs, #M0202L), which enabled insertion of the DD without introduction of any additional nucleotides or linker sequences. Plasmids were amplified by transforming XL-10 gold ultracompetent cells per the manufacturer’s protocol (Agilent Technologies, #200315), and final sequences were validated via DNA sequencing from Elim Biopharmaceuticals.

### Virus production

For third generation, self-inactivating lentiviral production, 15 million 293T cells were plated on a 15cm dish for 24 hours prior to transfection. On the day of transfection, 120uL of lipofectamine 2000 (Thermofisher, #11668500) was mixed with 1.5mL of room temperature Opti MEM (Fisher Scientific, #31-985-088) for 5 minutes. A mixture of 18ug pRSV-Rev plasmid, 18ug pMDLg/pRRe (Gag/Pol), 7ug pMD2.G (VSVG envelope), and 15ug pELNS vector plasmid in 1.5mL Opti MEM were added in dropwise fashion to the lipofectamine mixture and allowed to incubate for 30 minutes at room temperature. The 3mL transfection mixture was then added to 17mL of 293T complete medium (DMEM, 10% FBS, 1% Penicillin/streptomycin/glutamine), which was then applied to 293T cell plates. At 24-and 48-hours post-transfection, supernatant was collected, spun down at 500g, passed through at 0.45um syringe filter (Millipore, #SLHV033RB), and frozen at −80C. Retrovirus was produced in the 293GP packaging cell line as previously described (Long et al., 2015). Briefly, 70% confluent 293GP 15cm plates were co-transfected with 20ug MSGV vector plasmid and 10ug RD114 envelope plasmid with lipofectamine 2000. Viral supernatants were collected at 48-and 72-hours post-transfection, centrifuged to remove cell debris, and frozen at −80C for future use.

### T cell isolation

Primary human T cells were isolated from healthy donor buffy coats using the RosetteSep Human T cell Enrichment kit (Stem Cell Technologies), Lymphoprep density gradient medium, and SepMate-50 tubes according to the manufacturer’s protocol. Isolated T cells were cryopreserved at 1-2×10^7^ T cells per vial in CryoStor CS10 cryopreservation medium (Stem Cell Technologies).

### Lentiviral and retroviral transduction

Human T cells were transduced with lentivirus by adding thawed lentiviral supernatant directly onto 0.5-1×10^6^ T cells plated in 12-or 24-well plates. For retroviral transduction, non-tissue culture treated 12-or 24-well plates were coated overnight with 1mL or 0.5 mL of 25ug/mL retronectin in PBS (Takara), respectively. Plates were washed 2X with PBS and blocked with 2% BSA + PBS for 15 minutes. Thawed retroviral supernatant was added at ∼1mL per well and centrifuged at 3200 RPM for 1.5-2 hours at 32C before the addition of 0.5-1×10^6^ T cells.

### CAR-T cell culture

Cryopreserved T cells were thawed and activated on day 0 with Human T-Expander *α*CD3/CD28 Dynabeads (Gibco) at 3:1 bead:cell ratio in T cell media (AIMV supplemented with 5% FBS, 10mM HEPES, 2mM GlutaMAX, 100 U/mL penicillin, and 100ug/mL streptomycin (Gibco). Recombinant human IL-2 (Peprotech) was provided at 100 U/mL. Lentiviral transduction was done on day 1 and retroviral transductions were done on days 2 and 3 post-bead activation. Magnetic beads were removed on day 4, and T cells were fed with fresh culture medium every 2-3 days and normalized to 0.5 x 10^6^ cells/mL. For culture conditions requiring *α*PD-1, 5ug/mL nivolumab was added on D7 and supplemented every 2-3 days. For culture conditions requiring shield-1 (1uM unless otherwise noted), trimethoprim (1uM unless otherwise noted), dasatinib (1uM), or tazemetostat (1uM), freshly thawed drug was mixed with T cell culture medium and supplemented every 2-3d ays. HA.28*ζ*.FKBP CAR-T cells were rested via removal of either shield-1 or trimethoprim from the culture medium, whereby cells were washed 2X in PBS (changing conicals in between washes to ensure complete drug washout) and resuspended in fresh T cell culture medium lacking shield-1/trimethoprim.

### Immunoblotting

Whole-cell protein lysates were obtained in non-denaturing buffer (150 mmol/L NaCl, 50 mmol/L Tris-pH8, 1% NP-10, 0.25% sodium deoxycholate). Protein concentrations were estimated by using with DC Protein colorimetric assay (BioRad, 5000116). Per sample, 20μg of protein was mixed with 5X reducing loading buffer (Pierce, 39000), boiled at 95C for 5 min and loaded onto 11% PAGE gels. After electrophoresis, protein was transferred to PVF membranes. Signals were detected by enhanced chemiluminescence (Pierce) or with the Odyssey imaging system. Representative blots are shown. The following primary antibodies used were purchased from Cell Signaling: total ERK1/2 (no. 9102) and Phospho-ERK1/2 (no. 9101). The CD3*ζ* (4A12-F6) and phospho-CD3*ζ* (EP265(2)Y) antibodies were purchased from Abcam.

### Cytokine secretion assays

5×10^4^ CAR-T cells and 5×10^4^ tumor cells were cultured in 200uL T cell media (no supplemented IL-2) with or without shield-1/trimethoprim in 96-well flat bottom plates for 24 hours. For 1A7 anti-idiotype stimulation, serial dilutions of 1A7 were immobilized to Nunc Maxisorp 96-well ELISA plates (Thermo Scientific) in 1X Coating Buffer (BioLegend) overnight at 4C. Wells were washed once with PBS and 1×10^5^ CAR-T cells were plated in 200uL T cell media (no supplemented IL-2) and cultured for 24h. Triplicate wells were plated for each condition. In some conditions, stabilizing drug (shield-1 or trimethoprim) was added to DD-CAR-T cells in order to normalize CAR expression across experimental conditions (Figure 5A). Culture supernatants were collected and analyzed for IL-2 and IFNy concentration via ELISA using the manufacturer’s protocol (BioLegend).

### Incucyte killing assay

5×10^4^ GFP+ tumor cells were co-cultured with CAR-T cells in triplicate at a 1:1, 1:2, 1:4, or 1:8 Effector:Target ratio in 200uL T cell media lacking IL-2 in 96-well flat bottom plates. Plates were imaged every 2-6 hours for up to 120 hours using the IncuCyte ZOOM Live-Cell analysis system (Essen Bioscience). 4 images per well at 10X zoom were collected at each time point. Total integrated GFP intensity per well or total GFP area (um^2^/well) was assessed as a quantitative measure of live, GFP+ Nalm6-GD2 or GFP+ 143B-GL tumor cells, respectively. Values were normalized to the t=0 measurement. Effector:Target (E:T) ratios are indicated in the Figure legends.

### Flow cytometry and cell sorting

CAR-T cells were washed 2X in FACS buffer (2% FBS + PBS) prior to cell surface staining with fluorescent-labeled antibodies. Cells were resuspended in 100uL of antibody mastermix and incubated at 4C for 30 minutes. Cells were washed 2X in FACS buffer and analyzed on a BD Fortessa running FACS Diva software or sorted at the Stanford Shared FACS Facility on a BD FACSAria running FACS Diva software. For experiments involving staining of intracellular transcription factors, cells were surface stained as described above, then fixed, permeabilized and labeled according to the FoxP3 Transcription Factor Staining Buffer Set manufacturer’s protocol (eBioscience, # 00-5523-00). The 1A7 anti-14G2a idiotype antibody used to detect surface CAR was obtained from NCI-Frederick, which was conjugated in-house with Dylight488 and/or 650 antibody labeling kits (Thermo Fisher).

### Intracellular cytokine staining

CAR-T cells and 143B-GL were plated at a 1:1 E:T ratio in T cell medium (no supplemented IL-2) containing 1X monensin (eBioscience) and 1uL/test CD107a antibody (Clone H4A3, BioLegend) for 5-6 hours. After incubation, intracellular cytokine staining was performed using the FoxP3 Transcription Factor Staining Buffer Set according to the manufacturer’s protocol.

### Murine blood and spleen analysis

To calculate circulating numbers of adoptively transferred human T cells, peripheral blood was sampled from live mice via retro-orbital blood collection under isoflurane anesthesia. 50µL blood was labeled with anti-CD45 (eBioscience), lysed using BD FACS Lysing Solution, and quantified using CountBright Absolute Counting beads (Thermo Fisher) on a BD Fortessa cytometer. For phenotypic analysis of blood and spleen, mice were euthanized and blood was harvested via cardiac puncture. Spleens were mechanically dissociated and washed 2X in PBS. Both blood and spleen were treated with ACK lysis buffer (Fisher Scientific, # 50-751-7469) for 5 minutes at room temperature. The reaction was quenched with FACS buffer, washed 2X, stained with anti-human antibodies (as described in “Flow Cytometry”) and analyzed on a BD Fortessa cytometer.

### Mass Cytometry

1-2 x 10^6^ T cells from each culture condition were washed 2X in PBS (Rockland). Cells were resuspended in 1mL of 250nM cisplatin + PBS (Fluidigm Catalog: 201064), which was used to assess cell viability. After a 3-minute incubation at room temperature, the cisplatin reaction was quenched with cell staining media (CSM, 1X PBS with 0.05% BSA and .02% sodium azide). Cells were washed 2X in PBS and incubated in CSM + anti-CCR7 and anti-CXCR3 antibodies for 30 minutes at room temperature. These cell surface proteins were undetectable when labeled post-fixation, and thus, required labeling at this step. After 2 washes, cells were then fixed with 1.6% paraformaldehyde + PBS for 10 min at room temperature, followed by 2 washes with 1X PBS. Samples were subsequently flash frozen on dry ice for further use. Upon thawing and washing in CSM, barcoding was performed using a palladium-based approach using the Cell-ID 20-Plex Pd Barcoding Kit manufacturer’s protocol (Fluidigm Corporation). Cell samples were mixed together and treated as one sample for all subsequent steps. Titrated amounts of each cell surface antibody were mixed, filtered through a 0.1um spin filter (Millipore), and added to the merged barcoded sample for 30 minutes at room temperature. Cells were then washed 2X with CSM and permeabilized by adding ice cold methanol for 10 minutes at 4C, followed by two washes with CSM. Finally, samples were resuspended in IR-intercalator (0.5uM iridium intercalator and 1% PFA in 1X PBS), washed 1X with CSM and 2X with ddH2O, and filtered through a 50um cell strainer (Thermofisher). Cells were then resuspended at 1×10^6^ cells/mL in ddH2O with 1x EQ four-element beads (Fluidigm Corporation, #201078). Cells were acquired on a Fluidigm Helios mass cytometer.

### ATAC-seq

CAR-T cells were sorted on CD8 (see Cell Sorting) and processed for ATAC-seq, which was carried out as previously reported (Corces et al., 2017). Briefly, 50,000 cells from each sample were sorted by FACS, centrifuged at 500g at 4°C, then resuspended in 50uL of cold ATAC-seq Resuspension Buffer (RSB) (10 mM Tris-HCl, 10 mM NaCl, 3mM MgCl2) supplemented with 0.1% NP-40,0.1% Tween-20, and 0.01% Digitonin. Samples were incubated on ice for 3 minutes, then washed with 1 mL RSB supplemented with 0.1% Tween-20. Nuclei were pelleted at 500g for 10 minutes at 4°C. The nuclei pellet was resuspended in 50 μL transposition mix (25 μL 2x TD buffer, 2.5 μL transposase (Illumina), 16.5 μL PBS, 0.5 μL 1% digitonin, 0.5 μL 10% Tween-20, 5 μL H2O) and incubated at 37°C for 30 minutes in a thermomixer set to 1000 RPM. DNA clean up was achieved using the Zymo DNA Clean and Concentrator-5 Kit (#D4014). Libraries were PCR-amplified using the NEBNext Hi-Fidelity PCR Master Mix and custom primers (IDT) as described previously (Buenrostro et al., 2013). Libraries were sufficiently amplified following 5 cycles of PCR, as indicated by qPCR fluorescence curves (Buenrostro et al., 2013). Libraries were cleaned and purified using the Zymo DNA Clean and Concentrator-5 Kit and subsequently quantified with the KAPA Library Quantification Kit (#KK4854). Libraries were sequenced on the Illumina NextSeq at the Stanford Functional Genomics Facility with paired-end 75bp reads.

## QUANTIFICATION AND STATISTICAL ANALYSES

### Mass cytometry data processing and analysis

Acquired data was normalized and debarcoded using MATLAB-based software (Finck et al., 2013; Zunder et al., 2015). FCS files were imported into Cytobank and traditional handing gating was used to select single, viable CAR+/CD8+ T cells. New FCS files were then exported and arcsinh-transformed with a co-factor of 5, after which 2,000 cells were randomly downsampled per FCS file (i.e. per condition/timepoint). Additional columns were added to each FCS file representing: day of collection (numeric value 7-11), timepoint (numeric value 1-5), and exhaustion and memory scores. Exhaustion score for each cell was calculated as mean expression of PD-1, TIM-3, LAG-3, 2B4, CTLA-4, BTLA, and CD39 normalized to mean expression of these markers in control HA.28*ζ*.FKBP T cells cultured in the absence of shield-1 at each time point. Similarly, memory score was calculated using CD45RA, IL-7R, CD27, CD197 markers. Sampled/edited FCS files from days 7-11 for each condition were concatenated, creating an Always ON and Rested_D7-11_ files each with 10,000 total events. To create force-directed layouts (FDLs), we used *Vortex* software (Samusik et al., 2016). Edges, a basis for spring-like attractive forces between single cells within the FDL, connected each cell to its 10 nearest neighbors and were limited to cells of adjacent timepoints. Likewise, cell dissimilarity, which acts as repulsive forces between single cells within the FDL, was calculated based on angular (cosine) distance in indicated dimensions (Figure S2B).

### Bulk RNA-seq analysis

Bulk RNA-seq was performed by BGI America (Cambridge, MA) using the BGISEQ-500 platform, single end 50bp-read length, at 30 x 10^6^ reads per sample. Paired-end reads were aligned and quantified using Kallisto (version 0.44) (Bray et al., 2016) index against hg38 reference genome. The Gencode transcript annotations (version 27) were used for genomic location of transcriptomic units. Reads aligning to annotated regions were summarized as counts using R package tximport (version 1.12.3). Differential expression analysis of sample comparisons across various timepoints were performed using DESeq2 (version 1.24.0) (Love et al.). FDR cutoff of 0.05 was used for gene selection. Principal component analysis was performed on reads counts processed using variance-stabilizing transformation build into the DESeq2 package. PCA plots were generated using ggplot2 (2.3.2.1) in R (3.5.1). Gene set enrichment analysis was performed using the GSEA software (Broad Institute) as described (Mootha et al., 2003; Subramanian et al., 2005). Raw RNA-seq FASTQ files were used as input to perform quantitation of TCR receptor profiling using MiXCR tool (Bolotin et al., 2017). Briefly, reads were aligned to the V, D, J and C genes, assembled, and clonotypes were computed after filtering and error correction steps. Clonotypes were further assessed using tcR, which enabled computation of clontype abundance and diversity estimations based on the CDR3 amino acid length and V gene usage (Nazarov et al., 2015).

### ATAC-seq data processing

ATAC-seq libraries were processed following the pepatac pipeline (http://code.databio.org/PEPATAC/) using default options and aligned to hg19. Briefly, fastq files are trimmed to remove Illumina Nextera adapter sequences, and pre-aligned to the mitochondrial genome to remove mitochondrial reads. Multi-mapping reads aligning to repetitive regions of the genome were also filtered. Bowtie2 was used to align to the hg19 genome. Samtools was used to identify uniquely aligned reads and Picard was used to remove duplicate reads. The resulting deduplicated aligned BAM file was used for downstream analysis. Peaks were called on individual samples by the pepatac pipeline using MACS2. A merged peak list was created as previously described(Corces et al., 2018). All individually-called peaks were compiled, then rank-ordered by signal intensity (i.e., peak height). Starting with the first peak, any overlapping peaks are removed; any remaining peaks then overlapping with the second-strongest peak are removed, and so on, until a set of unique non-overlapping peaks remained. These peaks were resized to a standard 500bp width around the peak summit. Read counts in the merged peak set for each sample were generated with the bedtools intersect -c command. Merged signal tracks were made using aligned BAM files as inputs with the UCSC bigWigMerge and bedGraphToBigWig tools. The outlier samples (one D7 Always ON sample and one D15 Always OFF sample) were excluded from all ATAC-seq analyses, including merged conditions, as they did not cluster with replicates (see below ATAC-seq analysis).

### ATAC-seq analysis

The peak count table was pre-processed in R. It was depth normalized to account for differences in sequencing depth, then log-transformed with a pseudocount of 1. The resulting matrix was then quantile normalized across samples using the normalize.quantiles function in the preprocessCore library. Batch effects by patient were corrected with the removeBatchEffect function in the limma library. Differential peak calls across different conditions, using samples from different patients as replicates, were made using edgeR. P-values were adjusted by the Benjamini-Hochberg method. The number of significantly differentially accessible peaks was shown using the adjusted p-value less than or equal to 0.05 as a cutoff, aggregated by their logFC from the edgeR output. Samples were initially analyzed based on their pairwise Pearson correlation to determine if replicates clustered as expected. Only peaks with a read in at least half (18/36) of the samples were included, and the correlation was computed on the log-transformed counts. One outlier each from the D7 Always ON and D15 Always OFF samples were observed and subsequently excluded from downstream analyses. PCA plots were generated using the decomposition. PCA function in the sci-kit learn package in Python. PCA plots were generated using batch-corrected, quantile-normalized, log-transformed peak counts (with a pseudocount of 1). Only peaks with at least 5 counts-per-million in at least 9/36 of the samples were used. Homer was used to compute relative motif enrichment of the peaks contributing to the loadings of the first three components of the PCA analysis described above. The 5000 peaks most contributing to the PC in either direction based on their loadings were used. These peaks represent the peaks whose accessibility across samples most contributes to a given principal component. The findMotifsGenome function in Homer was used using these peaks relative to the background of the merged peak set (not the entire genome, but the merged peak set generated as described above). The motif enrichment thus represents the specific enrichment of those PCs against the background of CAR-T cell specific peaks, not just CAR-T cell specific peaks enriched over the entire genome. K-means clustering was performed on peaks using the cluster. KMeans function in sci-kit learn with the following arguments: n_cluster=6, n_init=100, max_iter=1000, precompute_distances=True. 6 clusters were chosen based on k=6 representing the inflection point of the sum of squared errors decline as k was increased from 1 to 25. K-means clustering was computed across the D15 samples, using only peaks that were significantly differentially expressed between any pair of D15 samples, z-normalized across samples. Heatmaps are shown clustered using ward clustering and Euclidean distance. ChromVAR was used to compute the relative differential accessibility of known transcription factor motifs across different samples as previously described (Rubin et al., 2019; Schep et al., 2017). Briefly, peaks counts were GC-corrected using the addGCBias function and the Jaspar motifs list was used to compute motif deviations with the getJasparMotifs, matchMotifs, and computeDeviations functions. Peak deviations were ranked by their variability across samples (standard deviation), and the most-variable deviations were shown. Note that some motifs are closely related and therefore behave similarly, and these closely related motifs were not collapsed or removed.

### Statistical Analysis

Unless otherwise stated, statistical analyses for significant differences between groups were conducted using 1-or 2-way ANOVA (unpaired, assumed Gaussian distribution, assumed equal standard deviations) and Dunnett’s multiple comparisons test (comparing all groups to Always ON or Vehicle controls) using GraphPad Prism 7. Survival curves were compared using the Log-rank Mantel-Cox test.

## ACKNOWLEDGEMENTS

This work was supported by the National Institutes of Health U54 CA232568-01 (C.L.M. and E.W.W.), RM1-007735 (H.Y.C.) and K08CA230188 (A.T.S.), a SU2C–St. Baldrick’s–National Cancer Institute Pediatric Cancer Dream Team Translational Research Grant (SU2CAACR-DT1113, C.L.M.) and the Parker Institute for Cancer Immunotherapy (A.T.S, H.Y.C, and C.L.M.). E.W.W. was supported by a Cellular and Molecular Immunobiology Training Grant (5 T32 AI07290, NIH NIAID, E.W.). H.Y.C. is an Investigator of the Howard Hughes Medical Institute A.T.S. was supported by a Bridge Scholar Award from the Parker Institute for Cancer Immunotherapy, a Career Award for Medical Scientists from the Burroughs Welcome Fund, and the Human Vaccines Project Michelson Prize for Human Immunology and Vaccine Research. We would like to acknowledge the National Cancer Institute (NCI) for providing the 1A7 anti-idiotype antibody.

## AUTHOR CONTRIBUTIONS

E.W.W. conceived of and designed this study, cloned the constructs, designed and performed experiments, analyzed data, and wrote the manuscript; R.C.L. and E.S. designed and performed experiments; E.S. performed immunoblot assays and generated GSEAs from RNA-seq data; J.L. and P.V. performed cell culture assays and in vivo experiments, ELISAs, and flow cytometry; P.X. mouse injections and bioluminescence imaging; M.M. performed mass cytometry experiments and designed and validated the antibody panel used in these experiments; Z.G. analyzed and interpreted mass cytometry data; H.A. and A.J.G. analyzed and interpreted RNA-seq data; H.A. Y.Q. and A.T.S. performed ATAC-seq experiments, and K.R.P., A.T.S., and H.Y.C. analyzed and interpreted ATAC-seq data; R.M. designed the HA.28*ζ* CAR and experiments; L.C. and T.J.W. assisted with the design and characterization of DD-CAR constructs; C.L.M. conceived of and designed this study, designed experiments, and wrote the manuscript; and all authors discussed the results and commented on the manuscript.

## SUPPLEMENTAL FIGURES

**Figure S1:**
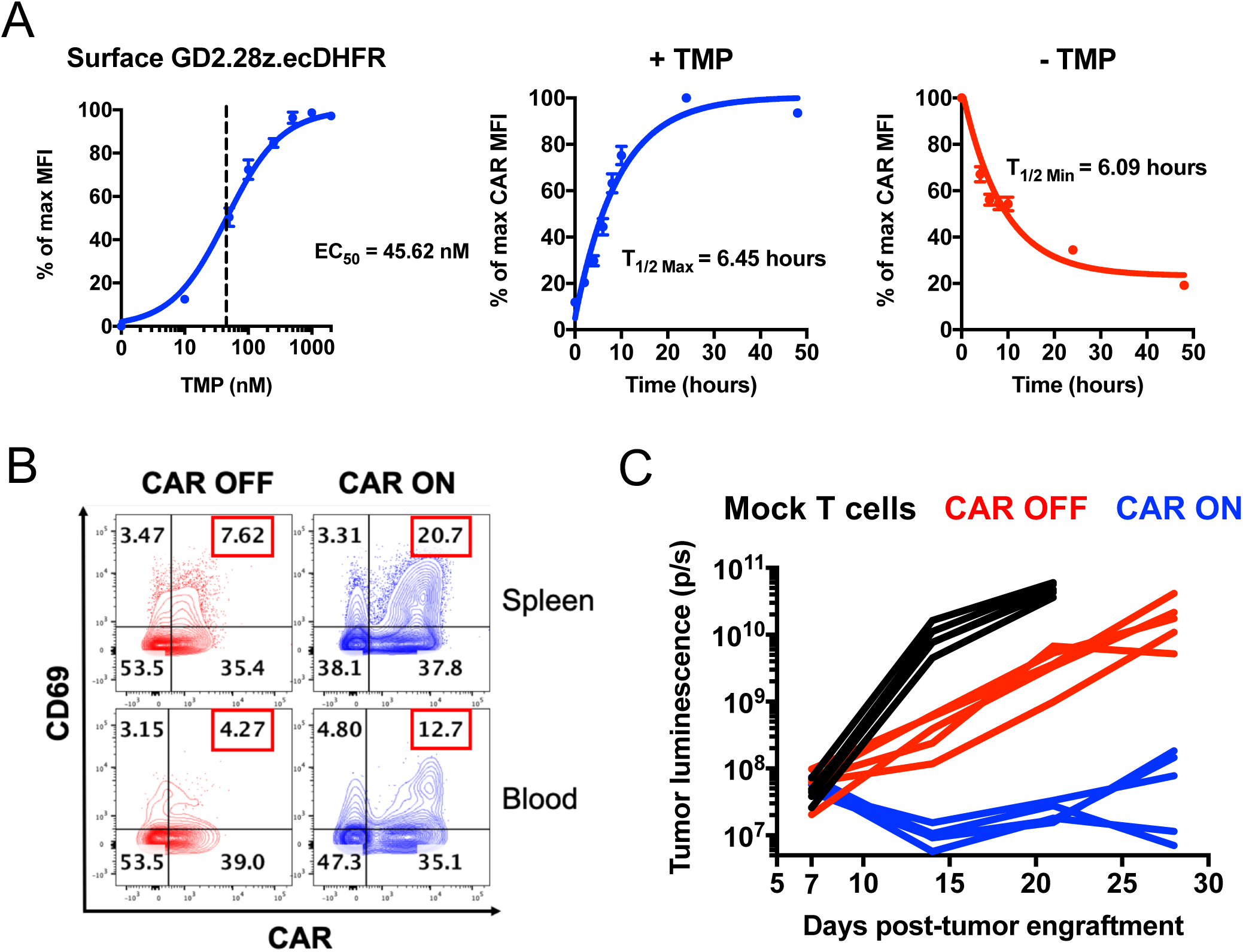
A GD2-targeting CAR fused to an ecDHFR destabilizing domain exhibits rapid ON/OFF kinetics and drug-dependent, analog control of expression and function *in vivo*. Related to Figure 1. **A)** CAR-T cells were stained with 1A7 anti-CAR idiotype antibody and analyzed via flow cytometry. Dose-dependent control (left) and ON/OFF kinetics (middle, right) were demonstrated via trimethoprim (TMP). CAR MFI was used to generate non-linear dose-response curves. Error bars represent mean ± SEM of 3 individual donors. **B)** 1×10^6^ Nalm6-GD2 leukemia cells were engrafted in mice. On days 5-7 post-engraftment, mice were dosed with vehicle (water, CAR OFF) or 200mg/kg trimethoprim (TMP, CAR ON) once daily. On day 7 post-engraftment, 1×10^7^ GD2.28*ζ*.ecDHFR CAR-T cells were infused. 24 hours after CAR-T cell infusion, peripheral blood and spleens were harvested for flow cytometry analyses. Contour plots demonstrate a TMP-dependent increase in co-expression of CAR and the activation marker CD69 in CAR ON mice compared CAR OFF mice. Representative mouse from n=7 total mice from 2 independent experiments. **C)** 1×10^6^ Nalm6-GD2 leukemia cells were engrafted in mice at day 0 and 2×10^6^ GD2.28*ζ*.ecDHFR CAR-T cells expanded *in vitro* in the absence of TMP for 15 days were infused IV on day 7 post-engraftment. Mice were dosed 6 days per week with vehicle (CAR OFF) or 200mg/kg TMP (CAR ON). Bioluminescence imaging of the tumor indicates TMP-dependent anti-tumor activity in CAR ON mice compared to CAR OFF mice (n=5 mice/group). Representative experiment (also shown in Figure 1F-G) from 3 individual experiments.

**Figure S2:**
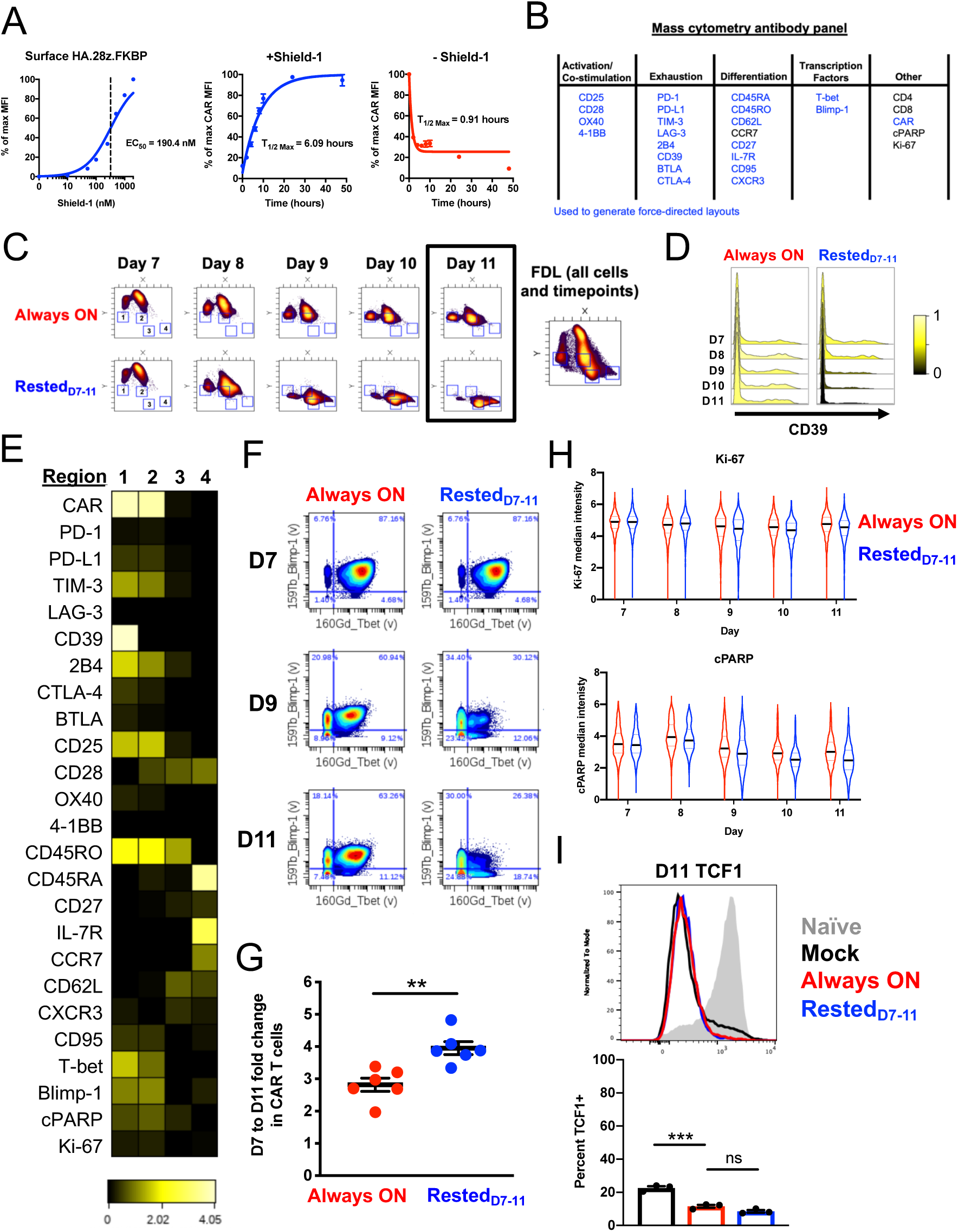
Mass cytometry demonstrates altered phenotypic features of rested HA.28ζ.FKBP CAR-T cells. Related to Figure 3. **A)** Flow cytometry of HA.28*ζ* CAR fused to an FKBP DD (HA.28*ζ*.FKBP) exhibits drug-dependent regulation of surface expression and rapid ON (T_1/2 Max_=6.09hr) and OFF kinetics (T_1/2 Min_=0.91hr). Error bars represent the mean ± SEM of 3 individual donors. **B)** Heavy metal-conjugated antibody panel used in mass cytometry experiments (27 antibodies, blue indicates markers used to generate the force directed layouts). **C)** Regions 1-4 (blue boxes) were identified based on the location of cells within the FDL on D11 (black box). **D)** CD39 expression over time in CD8+/CAR+ T cells. **E)** Heatmap demonstrates differences in cell phenotype between regions 1-4. Marker expression was normalized to the maximum value within each row. **F)** T-bet and Blimp-1 expression levels and co-expression frequency decreased over time in rested cells compared to Always ON cells. **G)** Fold change in CAR-T cell expansion between D7 and D11. Statistics were calculated using two-tailed paired student’s t test. **, p<0.01**. H)** Always ON and Rested_D7-11_ CAR-T cells exhibit similar levels of cleaved PARP-1 (cPARP) and Ki-67. Violin plots of single cells show quartiles with a band at the mean. **I)** Intracellular flow cytometry at D11 indicates low, comparable levels of TCF1 expression between Always ON and Rested_D7-11_ cells. TCF1+ gating was determined based on the TCF1+ naïve cell population. The histogram (top) shows data from 1 representative donor and the bar graph (bottom) represents the mean ± SEM of 3 individual donors. Statistics were calculated via one-way ANOVA and Dunnett’s multiple comparisons test. ***, p<0.001; ns, p>0.05. FDLs and other measurements from mass cytometry experiments were derived from 1 representative donor (n=3 individual donors).

**Figure S3:**
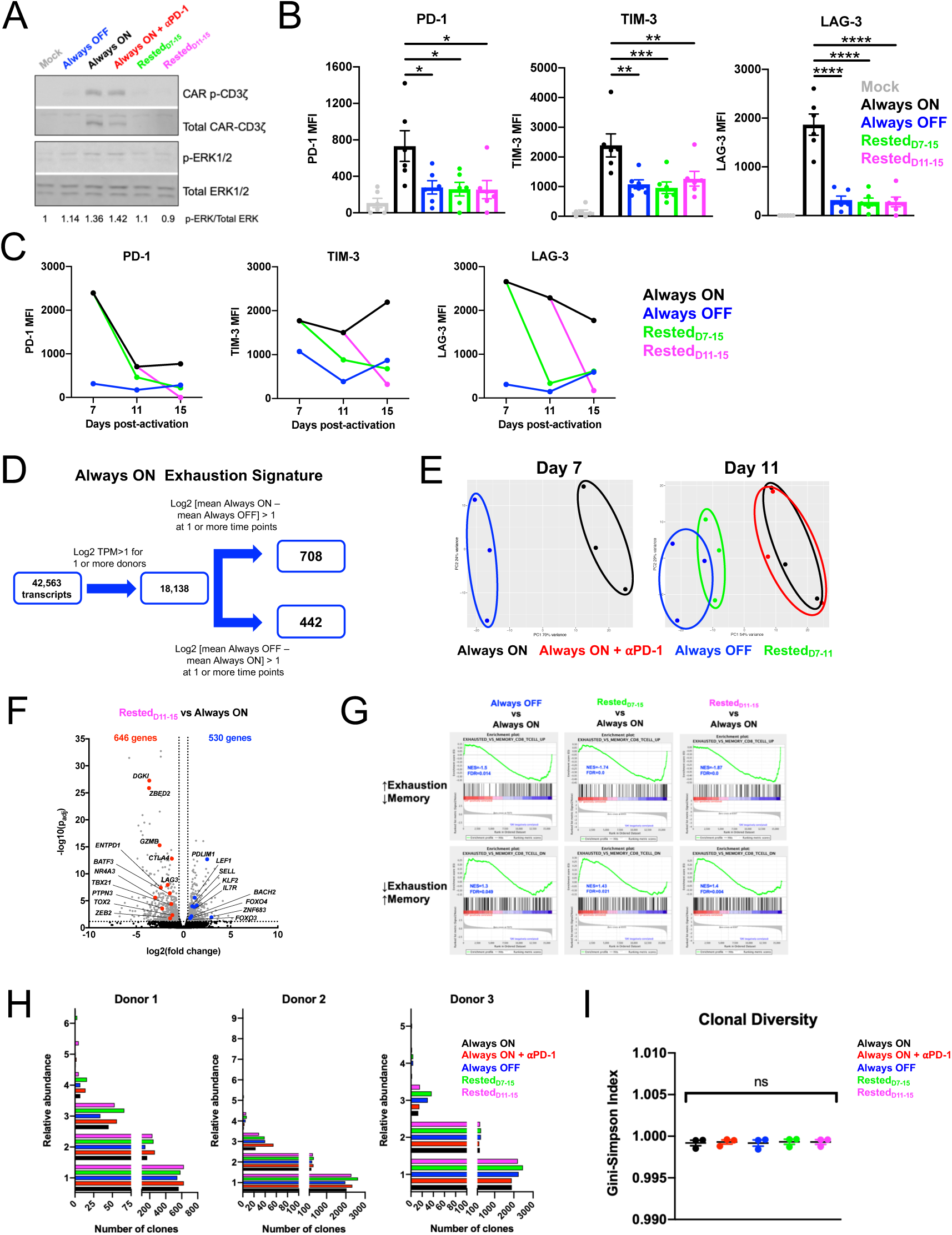
Phenotypic and transcriptional reprogramming of exhausted CAR-T cells via rest. Related to Figure 4. **A)** Western blot of CAR-T cells collected on D15 demonstrates reduced phosphorylated CAR CD3ζ and phosphorylated ERK1/2 in Rested_D7-15_ and Rested_D11-15_ CAR-T cells compared to Always ON cells. Representative donor from 3 individual donors. **B-C)** Flow cytometry demonstrates reversal of inhibitory receptor expression to Always OFF levels upon rest, regardless of whether rest is started at D7 (Rested_D7-11_, Rested_D7-15_) or D11 (Rested_D11-15_). (B) displays mean ± SEM of 6 individual donors and (C) displays one representative donor from 3 individual donors. **D-G)** Bulk RNA sequencing data of unsorted HA.28ζ.FKBP CAR-T cells (Figures 4E-G). **(D)** Data analysis pipeline for generating an Always ON exhaustion signature, which is comprised of 708 and 442 transcripts that are up-or downregulated in Always ON T cells, respectively. **E)** Principal component analyses of D7 (left) and D11 (right) RNA-sequencing data. **F)** Volcano plot of differentially expressed genes between Rested_D11-15_ cells and Always ON cells on D15. Genes were filtered for those that exhibited log2(fold change)>0.5 or <-0.5 and p(adjusted)<0.05. **G)** GSEA: gene sets that are downregulated in Always OFF, Rested_D7-15_ and Rested_D11-15_ cells compared to Always ON cells demonstrate significant overlap with genes upregulated in Exhausted versus Memory CD8+ (top) in a mouse model of chronic LCMV (Wherry et al. Immunity, 2007). Conversely, gene sets that are upregulated in Always OFF, Rested_D7-15_ and Rested_D11-15_ cells compared to Always ON cells demonstrate significant overlap with genes downregulated in Exhausted versus Memory CD8+ (top) in the LCMV model. GSEA – gene set enrichment analysis, NES – normalized enrichment score, FDR – false discovery rate. **H-I)** MiXCR analyses of TCR clonality on D15 RNA-seq samples show (H) the relative abundance of clones for each sample and (I) Gini-Simpson Index measuring clonotypic diversity for 3 donors. Bracketed bar in (I) indicates the same statistical significance for all sample comparisons. Statistics were calculated using one-way ANOVA and Dunnett’s (B) or Tukey’s (I) multiple comparison’s test. *, p<0.05; **, p<0.01 ***, p<0.001; ****p<0.0001; ns, p>0.05

**Figure S4:**
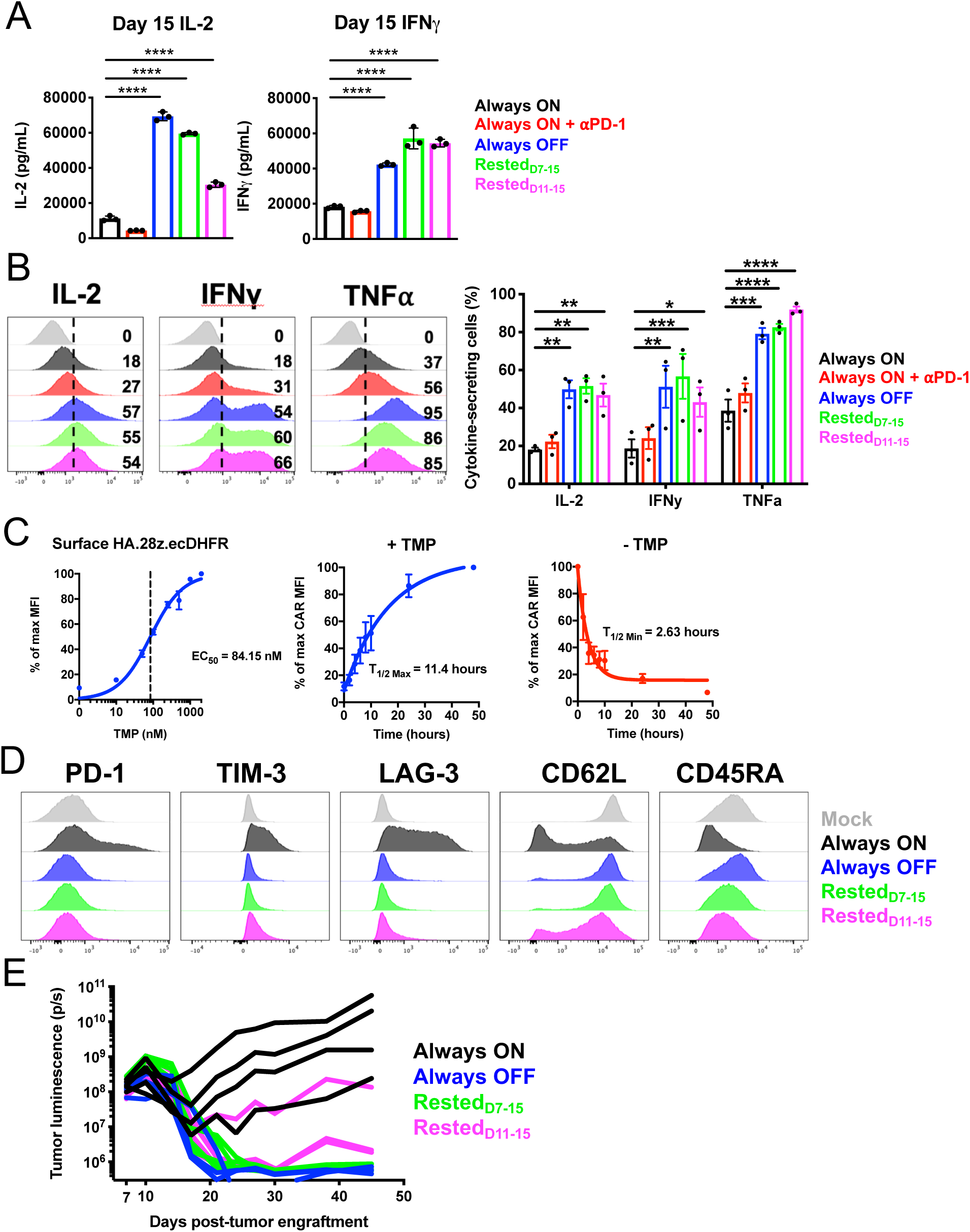
Transient rest functionally reinvigorates exhausted CAR-T cells. Related to Figure 5. **A)** HA.28ζ.FKBP CAR-T cell IL-2 (left) and IFNy (right) secretion in response to co-culture with Nalm6-GD2. Error bars represent ± SD of triplicate wells from one representative donor (related to Figure. 5C). **B)** Intracellular cytokine staining of CD8+ CAR+ T cells of rested groups demonstrates significantly increased frequency of cytokine-secreting cells. Representative histograms (left) and quantification of multiple donors (right) are shown. Error bars represent mean ± SEM of 3 individual donors. **C)** Dose-dependent control of HA.28ζ.ecDHFR expression (left) and ON/OFF kinetics (middle, right) were demonstrated via trimethoprim (TMP). CAR MFI was used to generate non-linear dose-response curves. Error bars represent mean ± SEM of 2-3 individual donors. **D)** Flow cytometry of HA.28ζ.ecDHFR conditioned as described in Fig. 4A. **E)** Quantification of tumor bioluminescence data from the experiments described in Figures 5H and 5I. Representative experiment from 3 individual experiments (n=3-5 mice/group). Statistics were calculated using twoway ANOVA and Dunnett’s multiple comparisons test. *, p<0.05; **, p<0.01 ***, p<0.001; ****p<0.0001

**Figure S5:**
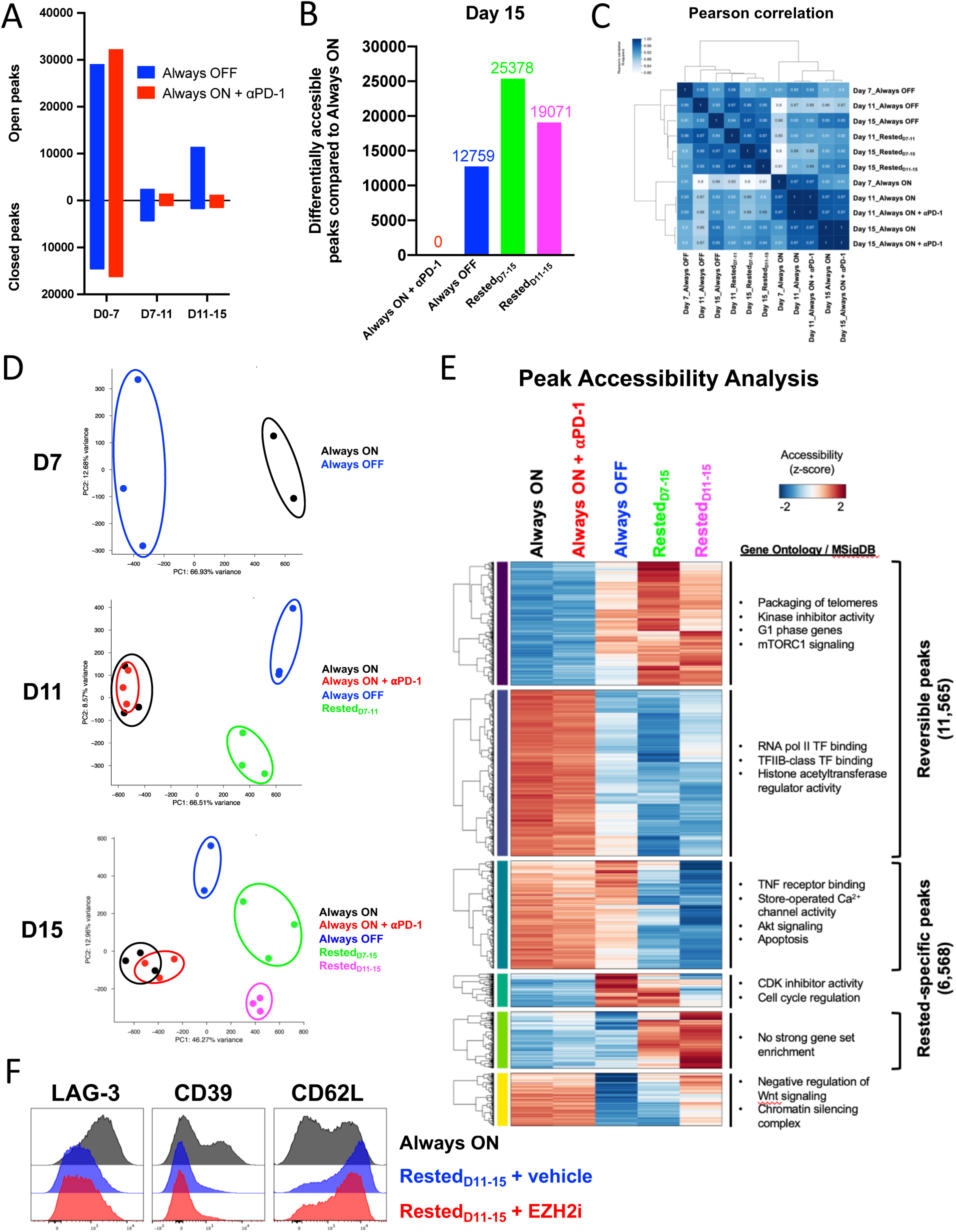
Epigenetic reprogramming of exhausted CAR-T cells via rest. Figure 6. **A)** Peak accessibility changes between timepoints were calculated based on p adjusted<0.05. Bar graphs display merged data from 2-3 donors. **B)** D15 differentially accessible peaks compared to Always ON (FDR<0.05). **C)** Pearson correlation and hierarchical clustering of ATAC-sequencing data from each condition and timepoint outlined in Figure 6A. Always OFF, Rested_D7-15_, and Rested_D11-15_ conditions cluster together, while Always ON and Always ON + aPD-1 comprise a different cluster. Heatmap generated with merged data from 2-3 individual donors. **D)** Unbiased principal component analysis of chromatin accessibility was assessed on D7, D11, and D15. Plots were generated with ATAC-seq data from 2-3 individual donors. **E)** K-means clustering of D15 differentially accessible peaks identified 6 distinct clusters. GSEA (gene ontology and MSigDB) of these clusters revealed biological processes that are enriched (packaging of telomeres, G1 phase) or diminished (Akt signaling, apoptosis, negative regulation of Wnt signaling, chromatin silencing complex) in rested conditions, consistent with quiescence and T cell memory. **F)** Flow cytometry at D15 demonstrates that tazemetostat treatment (EZH2i, red) does not affect phenotypic alterations that occur in Rested_D11-15_ cells compared to Always ON cells. Representative donor of 2 individual donors.

**Figure S6:**
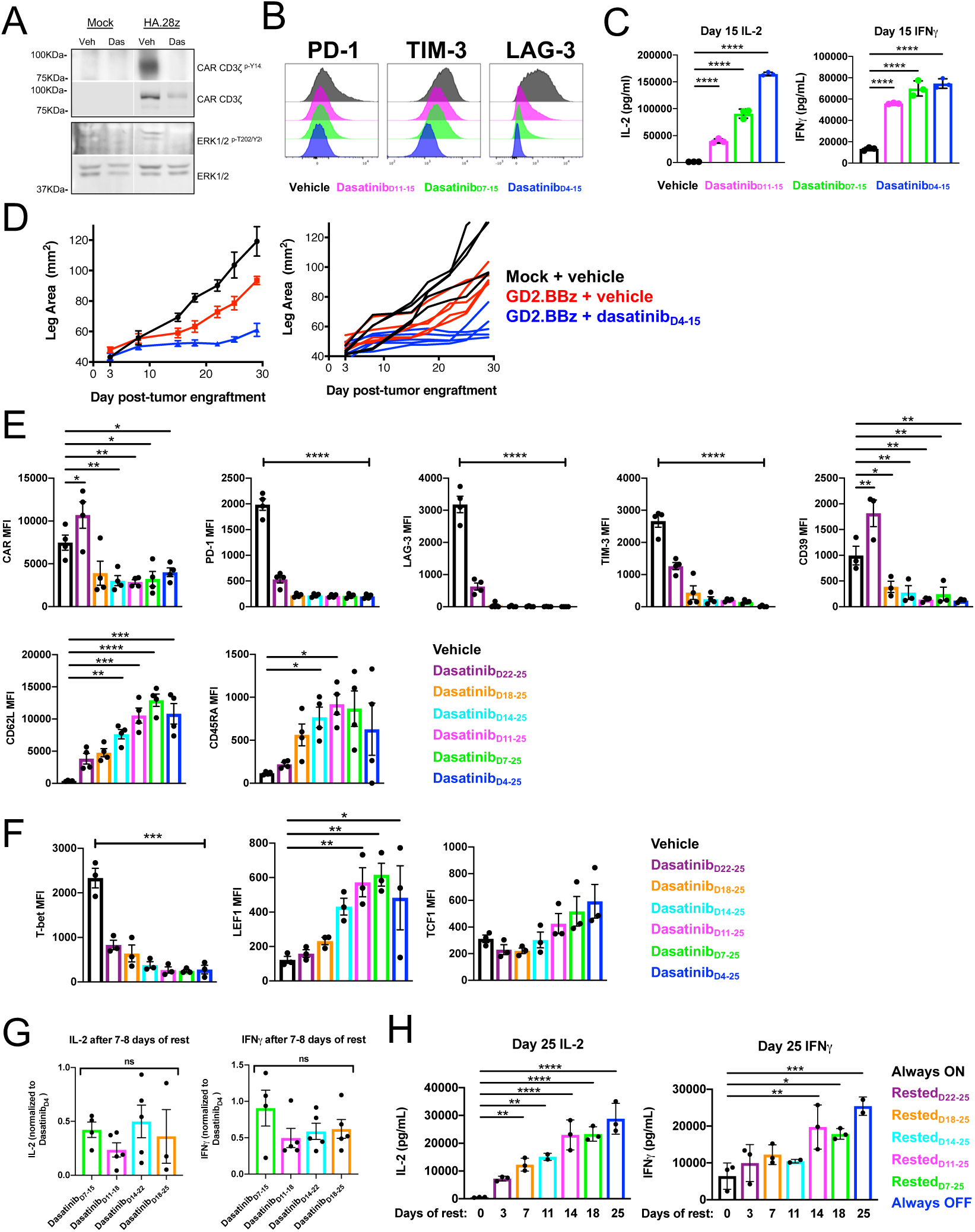
Dasatinib prevents tonic CAR signaling, reverses the exhaustion phenotype, and functionally reinvigorates exhausted CAR-T cells. Related to Figure 7. **A)** D15 HA.28*ζ* CAR-T cells cultured in the presence of dasatinib from D4-15 (DasatinibD4-15) demonstrates reduced phosphorylation of CAR CD3*ζ* and phosphorylation of ERK1/2 compared to vehicle-treated cells (Veh). Representative donor from 3 individual donors. **B)** Flow cytometry of D15 CD8+ CAR+ T cells demonstrates reversal of inhibitory receptor expression in cells rested via dasatinib (DasatinibD7-15, DasatinibD11-15) compared to exhausted vehicle-treated controls (Vehicle). Histograms show 1 representative donor of 3 individual donors. **C)** Dasatinib was removed from CAR-T cell cultures approximately 16 hours prior to co-culture with Nalm6-GD2 leukemia on D15 and IL-2 and IFNy secretion was subsequently assessed. Error bars represent mean ± SD of triplicate wells from 1 representative donor (n=3 donors). **D)** 0.5×10^6^ 143B-GL osteosarcoma cells were engrafted in the flank of NSG mice, and 1×10^7^ CAR+ GD2.BB*ζ* CAR-T cells that had been cultured in the presence of vehicle (DMSO) or dasatinib for 10 days were infused on day 3 post-engraftment. Tumor size was monitored via caliper measurements. CAR-T cells cultured in dasatinib exhibited superior tumor control to those cultured in vehicle. Error bars represent mean ± SEM of 5 individual mice from 1 experiment. **E-F)** Quantification of D25 **(E)** exhaustion marker and **(F)** transcription factor expression via flow cytometry. Error bars represent mean MFI ± SEM of 3-4 individual donors. Bracketed horizontal bars indicate the same statistical significance across all conditions compared to Vehicle controls. **G)** Co-culture assays with Nalm6-GD2 leukemia were conducted on D15, D18, D22, and D25. IL-2 and IFN*γ* secretion levels from CAR-T cells cultured with dasatinib for 7-8 days were normalized to DasatinibD4-25 for each individual timepoint. Graphs display data from 4-5 individual donors. These data are also represented in Figure 7H, but were replotted here to illustrate cytokine secretion when duration of rest is normalized across conditions. **H)** HA.28*ζ*.FKBP CAR-T cells were cultured in a protracted *in vitro* time course, as outlined in Fig. 7B. Similar to data shown with dasatinib (Fig. 7F-H), rested HA.28*ζ*.FKBP CAR-T cells exhibit reinvigoration at both early and late timepoints compared to Always ON cells. Furthermore, their capacity for reinvigoration is correlated to duration of rest. Error bars represent mean ± SD of 2-3 wells from one representative donor (n=3 individual donors). Statistics were calculated using one-way ANOVA and Dunnett’s multiple comparisons test. *, p<0.05; **, p<0.01 ***, p<0.001; ****p<0.0001; ns, p>0.05

